# RAD52 prevents accumulation of RAD51 and Polα-dependent DNA gaps at perturbed replication forks

**DOI:** 10.1101/2023.04.12.536536

**Authors:** Giorgia Marozzi, Ludovica Di Biagi, Eva Malacaria, Masayoshi Honda, Pasquale Valenzisi, Francesca Antonella Aiello, Maria Spies, Annapaola Franchitto, Pietro Pichierri

## Abstract

Replication gaps can arise as a consequence of perturbed DNA replication, and their accumulation might undermine the stability of the genome. Loss of RAD52, a protein involved in the regulation of fork reversal, promotes accumulation of parental ssDNA gaps during replication perturbation. Here, we demonstrate that this is due to the engagement of Polα downstream of the extensive degradation of perturbed replication forks after their reversal and is not dependent on PrimPol. Polα is hyper-recruited at parental ssDNA in the absence of RAD52, and this recruitment is dependent on fork reversal enzymes and RAD51. Of note, we report that the interaction between Polα and RAD51 is stimulated by RAD52 inhibition, and Polα-dependent gap accumulation requires formation of the RAD51 nucleoprotein filaments. Our data indicate that the RAD51/Polα-dependent repriming is essential to support fork progression, limit DNA damage and improve viability of RAD52-deficient cells when replication is perturbed. Altogether, this study shows that RAD51/Polα-dependent repriming is a genuine fork recovery mechanism activated to overcome loss of RAD52 function at replication forks.

## INTRODUCTION

Correct and timely response to perturbed replication forks prevents accumulation of unreplicated DNA regions, DNA damage and genomic rearrangements ^1–4^. For this reason, cells evolved multiple mechanisms of protection that deal with impaired replication fork progression and replication stress ^3,5,6^. A crucial step in this process is represented by the DNA remodelling occurring at perturbed or stalled replication forks. Fork remodelling involves the reannealing of the two nascent strands promoted by regression of the replication fork structures producing a four-way DNA intermediate called reversed fork (RF) ^7^. This reaction has been first discovered in bacteria where it is catalysed by RecG while, in human cells, involves multiple factors including SMARCAL1, ZRANB3, HLTF, FBH1 and the RAD51 recombinase ^6,8,9^. While contributing to protection of nascent ssDNA, fork reversal produces a DNA end that can be recognised and processed by nucleases. The RF extruded strand is protected by unscheduled and extensive degradation by BRCA2, RAD51 and other proteins acting as “barriers” against the action of MRE11 and EXO1 ^5,10–12^. Pathological DNA transactions at replication forks is also counteracted by proteins that regulate fork remodelling, such as ATR or RADX ^13,14^. In addition, our previous work uncovered a role for RAD52 in preventing fork degradation by regulating the SMARCAL1 access to the perturbed replication fork ^15,16^. Since the RF can be also a target for exo- or endonucleases ^3,5^, cells balance its usage with other mechanism granting continuation of DNA synthesis. Recently, it has been reported that replication fork reversal and repriming act as parallel “fork recovery” pathways: limited fork reversal upregulates repriming and vice versa ^17–19^. In yeast, repriming under perturbed replication involves the Polα primase but recent reports indicate that a specialised Primase-Polymerase, PrimPol is responsible for repriming in human cells ^20–23^.

Repriming at perturbed replication forks leads to accumulation of ssDNA gaps that needs to be subsequently repaired to prevent accumulation of unreplicated DNA in mitosis ^24,25^. Intriguingly, abrogation of RAD52 function stimulates formation of parental ssDNA although it does not overtly affect the ability of fork to restart after stalling ^15^.

Here, we used multiple cell biology techniques and cell models to investigate if loss of RAD52 function could affect the repriming in response to perturbed replication. We show that loss of RAD52 function stimulates the accumulation of parental ssDNA gaps and that those gaps are not dependent on PrimPol but rather are dependent on Polα. In the absence of a functional RAD52, origin-independent Polα recruitment at DNA is stimulated.

Such origin-independent Polα recruitment occurs downstream of fork reversal and degradation and is linked to inability to induce MUS81-dependent DSBs at the degraded RFs. Notably, the recruitment of Polα licensed by RAD52 inactivity requires RAD51 nucleoprotein filament formation and involves binding with RAD51 itself. Under these conditions, engagement of RAD51/Polα-mediated repriming ensures maximal fork progression and DNA damage avoidance during replication stress. However, this mechanism also results in ssDNA gaps left behind the fork. Therefore, our results uncover a novel mechanism of repriming granting fork recovery and protecting from genome instability engaged when aberrant or partial DNA transactions occur at perturbed replication forks.

## RESULTS

### RAD52 prevents formation of gaps at perturbed replication forks

Perturbed replication forks are extensively degraded in the absence of RAD52; however, a large part of them retains the ability to resume DNA synthesis ^15^. Two main pathways are used to resume degraded DNA replication forks: recombination and repriming ^3,25,26^. Since RAD52 inactivation prevents DSBs formation at degraded forks and subsequent break-induced replication ^11^, we analysed whether replication fork recovery involved repriming events in the absence of active RAD52. To this end, we monitored the exposure of parental ssDNA by a native IdU detection ^27^. As depicted in the cartoon of Figure 1A, parental ssDNA may represent DNA gaps left behind the replication fork or can arise downstream extensive nucleolytic degradation at the fork or after its reversal. Parental (template) DNA was labelled with Iododeoxyuridine (IdU) for 24h followed by a 2h chase to prevent cross-labelling of nascent DNA prior to treatment with HU in the presence or not of the RAD52 inhibitor ECG (RAD52i; ^28^) (Figure 1A). Our previous data indicate that inhibition of RAD52 did not result in the accumulation of parental ssDNA during treatment with 2mM HU ^15^. Thus, parental ssDNA was examined at various recovery times post-HU (Figure 1A). Parental ssDNA was detected in both RAD52-proficient and RAD52-deficient cells after 2h of recovery. However, it was significantly higher in RAD52-inhibited cells after 4h and persisted at 18h, when it still exceeded the amount observed in the non-inhibited cells.

**Figure 1.**
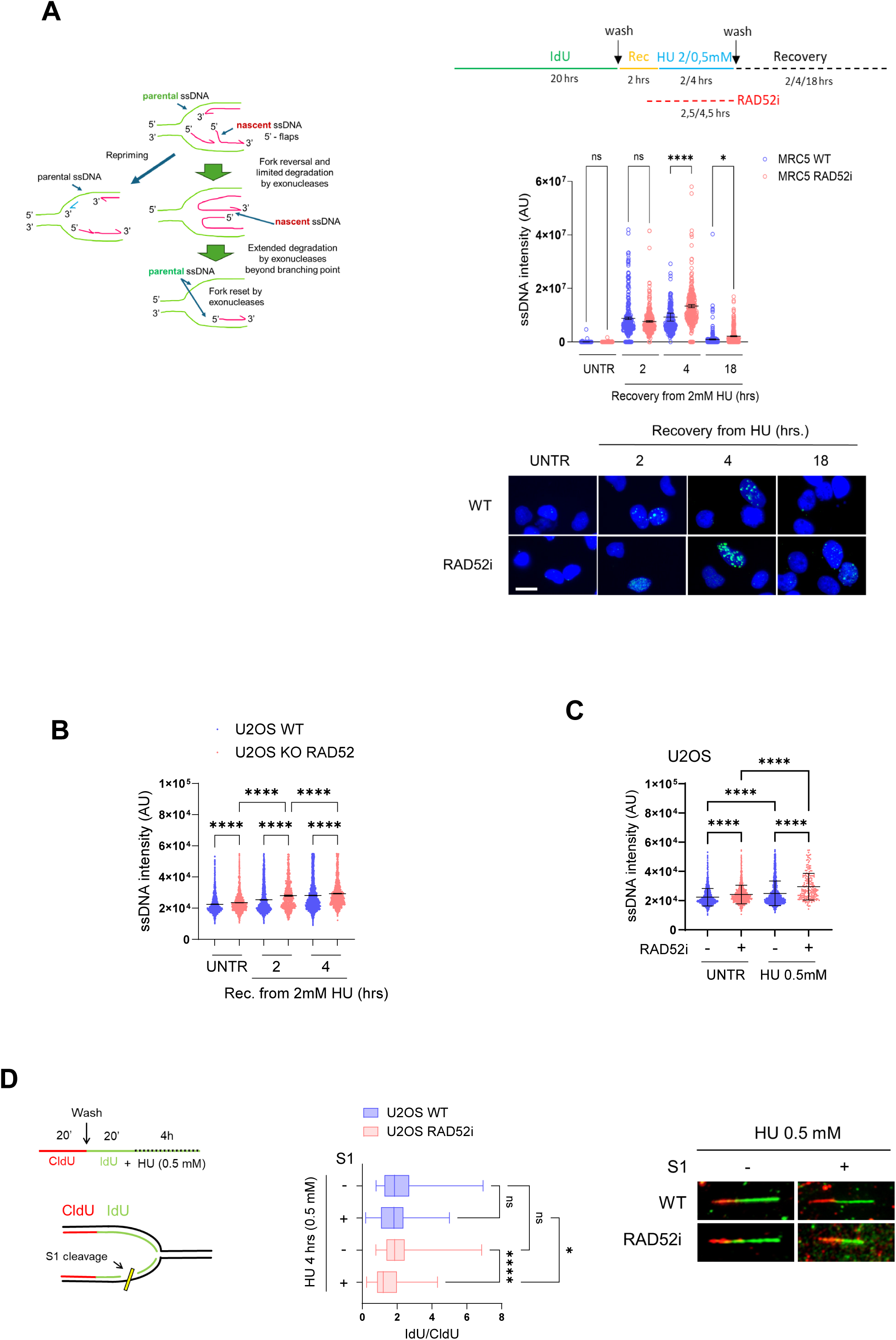
RAD52 deficiency stimulates repriming. **A.** Analysis of parental ssDNA exposure by immunofluorescence. MRC5 WT were treated as indicated on the experimental scheme. In untreated cells, RAD52i (50μM) was left for 4 hours. The ssDNA was detected by native anti-IdU immunofluorescence. Graph shows the intensity of ssDNA staining per cell from three replicates. Representative images are shown. Scale bar represents 10 µm.**B and C**) QIBC analysis of parental ssDNA by immunofluorescence. Parental or RAD52 KO cells were treated as above. Cells were acquired and analysed by Olympus ScanR High Content Imaging System. At least 1600 cells were analysed for each point from three replicates. **D**. Analysis of ssDNA gaps using S1-fiber assay. Cells were labelled as indicated in the scheme and treated or not with the S1 nuclease before spreading the DNA. The graph shows the IdU/CldU tract length ratio. Shorter ratios indicate ssDNA gaps. Representative images of DNA fibres are presented. (ns = not significant; *P < 0.1; **P < 0.01; ***P<0.001; ****P < 0.0001; Kruskall-Wallis test (B-D) or Mann–Whitney test (A).

To further confirm that inhibition of RAD52 leads to accumulation of parental ssDNA in response to stalled or perturbed replication, we performed Quantitative Image-Based Cytometry (QIBC) using RAD52 KO cells. Quantitative imaging confirmed the increased detection of parental ssDNA in cells inhibited of RAD52 and showed a significant increase of ssDNA in S-phase as compared with the non-inhibited cells, while no statistically significant differences were observed in G2 (Figure 1B and Supplementary 1A). Such elevated parental ssDNA is unrelated to the resection of DNA ends after breakage, at least at the earlier recovery times, since loss of RAD52 prevents DSBs at perturbed forks ^15^.

Similar to the results with RAD52i and KO cells, higher levels of parental ssDNA were observed during replication recovery in cells with stable shRAD52 ^15^ expression at 4 and 18h of recovery, and in U2OS cells (Supplementary Figure 1B and C). In U2OS cells, RAD52-inhibition stimulated the accumulation of parental ssDNA at 2h and 4h of recovery after 2mM HU even if differences were not significant at 4h of recovery (Supplementary Figure 1C). Loss of RAD52 simulates MRE11-mediated fork degradation and ssDNA formation at perturbed forks ^15^, however, MRE11 inhibition did not have much effect on parental ssDNA levels in RAD52-deficient cells at 2h of recovery, although it reduced the level of parental ssDNA at 4h during recovery from 2mM HU (Supplementary Figure 1C). Subsequently, we analysed parental ssDNA in U2OS cells treated with 0.5mM HU, a dose that slows replication without causing complete stalling ^29^ and stimulates parental ssDNA exposure. Although treatment with 4h of 0.5mM HU exposed parental ssDNA in U2OS cells, the detection of parental ssDNA significantly increased when RAD52 was inhibited (Figure 1C). Treatment with MIRIN did not significantly affect parental ssDNA exposure at 2h of treatment with 0.5mM HU even if a minor, but not significant, reduction was observed at 4h of treatment (Supplementary Figure 1D). Neither acute RAD52 inhibition nor RNAi knockdown altered S-phase cell proportions or IdU incorporation, excluding these factors as explanations for the observed effects (Supplementary Figure 2). As previously shown ^15^, RAD52 inhibition did not affect fork progression rates per se in untreated cells (Supplementary Figure 3). Hence, to demonstrate that the elevated levels of parental ssDNA detected in the absence of RAD52 correlated with the presence of replication-dependent DNA gaps, we performed the S1 DNA fiber assay ^30^. We pulse-labelled cells with CldU followed by labelling with IdU and treatment with 0.5mM HU, in the presence or not of the RAD52i (see scheme in Figure 1D). After treatment, cells were exposed to S1 nuclease to cut regions of ssDNA before obtaining DNA fibers. Intact replication tracts typically show an IdU/CldU ratio > 1, since IdU is incorporated, albeit slowly, during the 0.5 mM HU treatment. In contrast, gap-containing tracts become shorter after S1 nuclease treatment, resulting in an IdU/CldU ratio < 1. After 0.5mM HU treatment, RAD52-inhibited cells showed an IdU/CldU ratio similar to control cells (Figure 1D; -S1). However, S1 nuclease reduced the ratio significantly in RAD52-inhibited cells, suggesting RAD52 deficiency leads to daughter-strand DNA gaps during impaired replication.

These findings indicate that inhibition or depletion of RAD52 results in the accumulation of parental ssDNA correlating with a stimulated formation of DNA gaps in response to perturbed replication.

### Inhibition of RAD52 stimulates Polα-dependent gaps

The accumulation of gaps at perturbed replication forks in BRCA-deficient cells or in some other pathological conditions attributed to a defect in the metabolism of Okazaki fragments is correlated with PrimPol-mediated repriming ^17,31,32^. Thus, we tested if inhibition of RAD52 similarly stimulates the repriming activity of PrimPol. To analyse gaps, we performed the S1 DNA fiber assay in wild-type or PrimPol KO MRC5SV40 cells ^33^, treated with 0.5 mM HU in the presence or absence of RAD52i. As expected, wild-type cells (WT) did not show a significant accumulation of DNA gaps in the absence of RAD52i (Figure 2A, compare ±S1). PrimPol KO cells showed relatively longer IdU tracts in 0.5mM HU respect to WT cells. However, the length of these IdU tracts was minimally affected by S1 (Figure 2A). Inhibition of RAD52 in WT cells confirmed the formation of DNA gaps following replication perturbation. Indeed, S1 treatment reduced the IdU tract-length and resulted in IdU/CldU ratios that were lower than those without prior S1 treatment (Figure 2A). Interestingly, inhibition of RAD52 stimulated teh formation of DNA gaps also in PrimPol KO cells since the IdU/CldU ratios were reduced by S1 treatment (Figure 2A).

**Figure 2.**
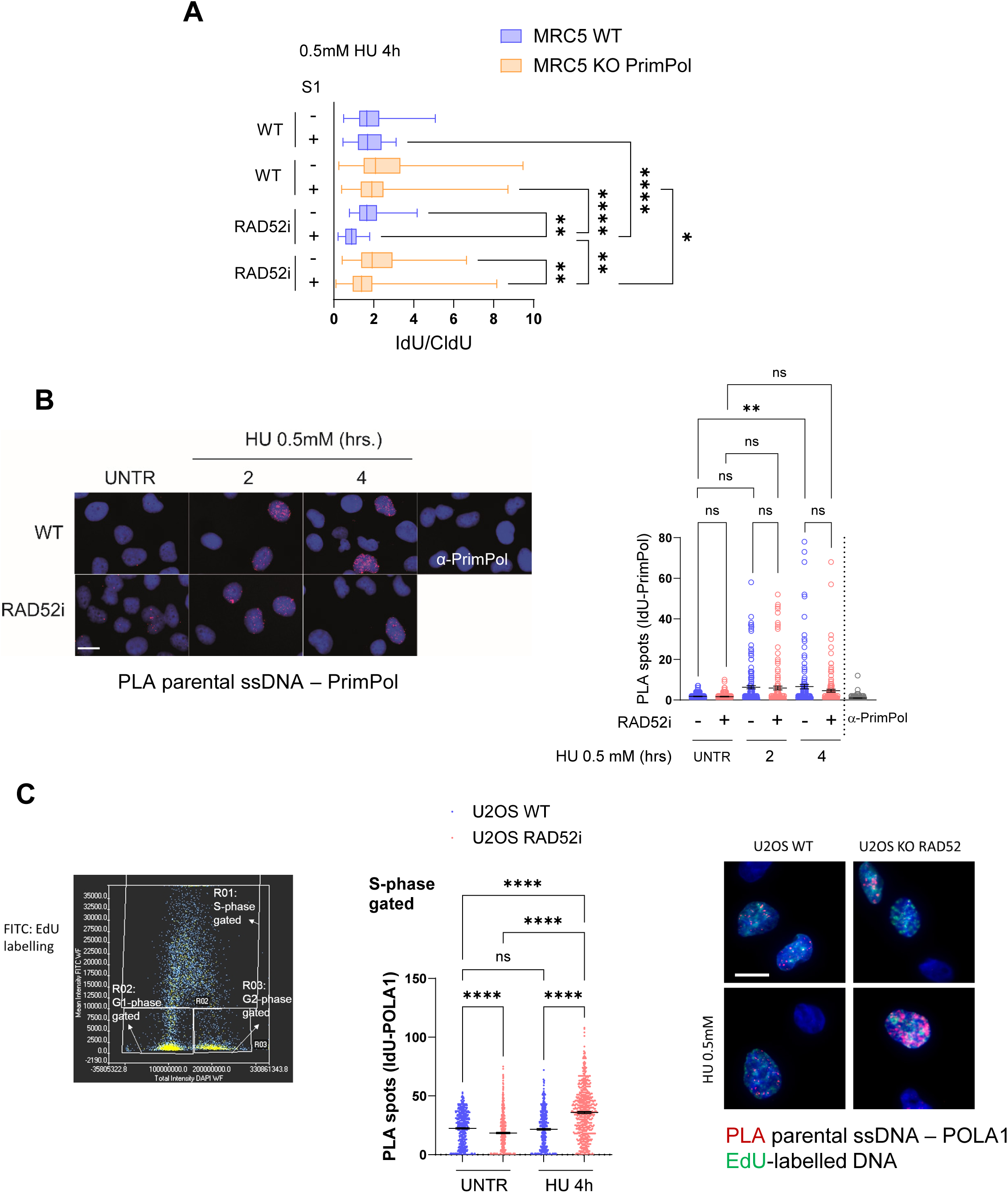
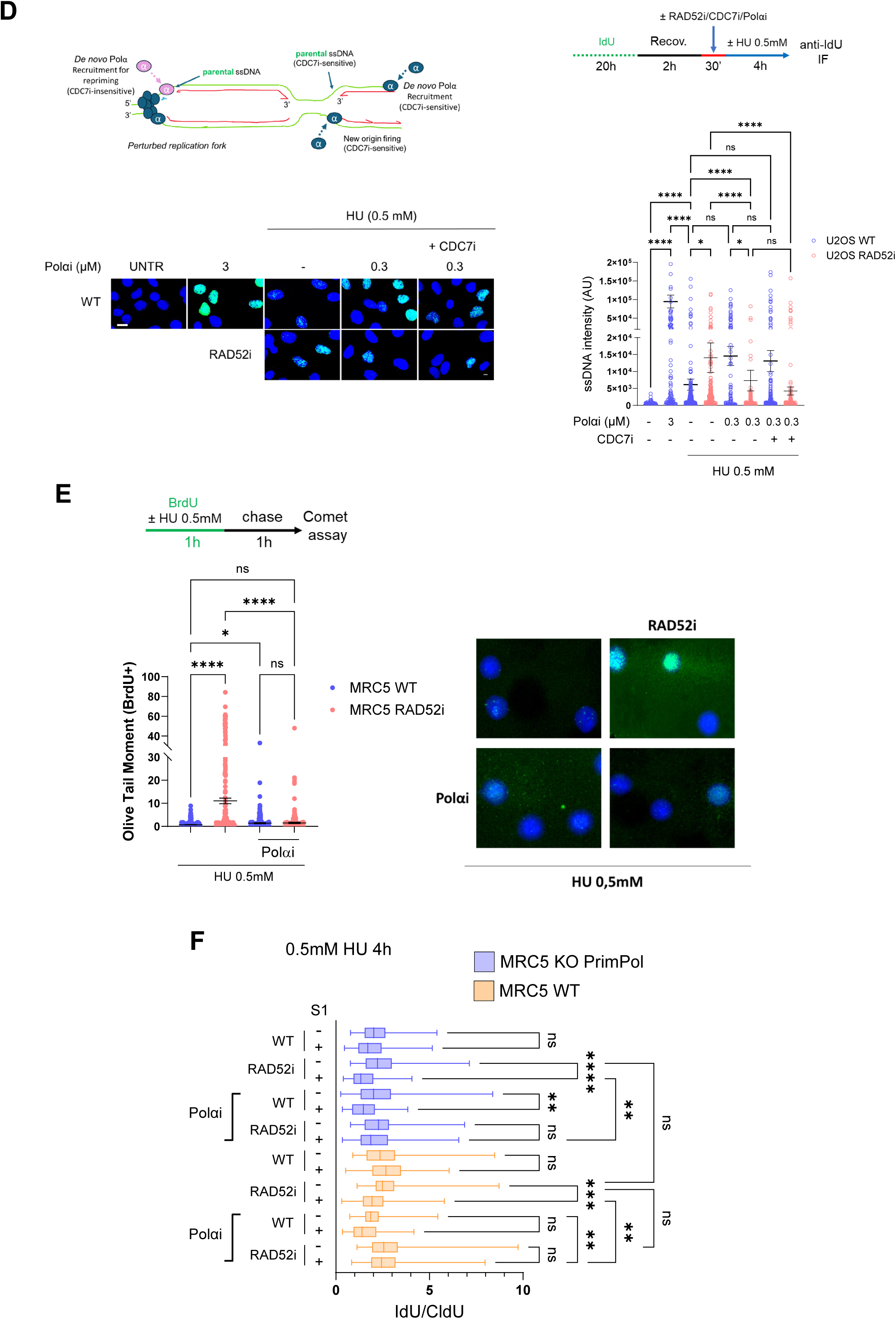
Parental DNA gaps are Polα-, but not PrimPol-, dependent in the absence of RAD52. **A.** Analysis of ssDNA gaps through S1 fiber assay. Wild-type MRC5 or KO PrimPol were treated with 0.5mM HU for 4 hours. RAD52i was given 30 min before replicative stress induction. The graph reports the mean IdU/CldU ratio. (*P < 0.1; **P < 0.01; ****P < 0.0001, Kruskall-Wallis test. Where not indicated, samples are not statistically significant). **B.** Analysis of PrimPol-ssDNA interaction using Proximity Ligation Assay (PLA). The PLA reaction was carried out using antibodies against PrimPol and IdU. Controls were subjected to PLA with anti-IdU or anti-PrimPol only. The graph reports the number of PLA spot per nucleus. Representative images are shown (ns = not significant; **P<0.1; Kruskal-Wallis test). **C.** Analysis of ssDNA-Polα interaction using PLA. Cells were treated with Hu 0.5mM for 4h. The images were acquired and analysed by Olympus ScanR High Content Imaging System. Cells were gated according to the cell cycle phase that was identified by plotting the total intensity of DAPI (x-axis) on the mean intensity of EdU labelling (y-axis): actively replicating cells will be positive for the EdU signal. **D.** Effect of mild Polα inhibition on parental ssDNA exposure. in U2OS treated with to. Cells were treated as indicated on the experimental scheme; CDC7i was used to block new origin firing. Graph shows the ssDNA staining (AU) per cell (ns = not significant; *P < 0.5; ****P < 0.0001; Kruskal–Wallis test). Representative images are shown. **E**) BrdU alkaline Comet assay to show that Polαi suppress DNA gaps in RAD52i cells. Nascent DNA is labelled with BrdU as explained in the scheme. The graph shows individual Olive Tail Moment values in BrdU+ nuclei. Representative images are shown. Means ± SE are presented for each sample (ns = not significant; *P < 0.1; ****P < 0.0001, Kruskal-Wallis test). **F.** Analysis of ssDNA gaps through S1 fiber assay. Wild-type MRC5 or MRC5 KO PrimPol were treated with Polαi (0.3 μM) and RAD52i 30 min before HU (0.5 mM; 4h). The graph shows the IdU/CldU ratio. (ns = not significant; *P < 0.5; **P<0.1; ***P < 0.001; ****P < 0.0001; Kruskal–Wallis test).

Next, we decided to investigate the recruitment of PrimPol at parental ssDNA following fork arrest by *in situ* PLA ^15,27^. PrimPol did not appear to be involved in DNA gap formation when RAD52 is inhibited (Figure 2A). Although PrimPol recruitment to parental ssDNA increased after 0.5mM HU, there was no such enhancement in RAD52-inhibited cells compared to wild-type (Figure 2B). Similar results were obtained in PrimPol KO cells complemented with GFP-PrimPol (Supplementary Figure 4).

Gap formation occurred independently of PrimPol and no increased recruitment of PrimPol was observed after replication perturbation in RAD52-inhibited cells, suggesting that other proteins might play a role in repriming events and prompting us to investigate alternative candidates. Therefore, we investigated whether inhibition of RAD52 affected the recruitment of Polα. Thus, we performed a parental ssDNA PLA with an antibody against the POLA1 subunit. Analysis of PLA was performed by QIBC in S-phase gated cells, as shown in the exemplification of plot of Figure 2C, and spots were counted accordingly using a ScanR system. The quantitative analysis of the parental ssDNA-POLA1 PLA spots evidenced the presence of Polα already in untreated cells, which is consistent with the generic role during DNA replication. Treatment with 0.5mM HU did not increase the number of PLA spots in wild-type cells but the presence of POLA1 at parental ssDNA was significantly more elevated in cells treated with the RAD52 inhibitor (Figure 2C). Combining parental ssDNA-POLA1 PLA with EdU immunofluorescence to label S-phase cells, we confirmed that EdU-negative cells displayed only few PLA spots – typically seven or less – and this unexpected localisation was not affected by RAD52 inhibition, although appeared to be more represented in HU-treated cells (Supplementary Figure 5A, B). Although Inhibition of RAD52 stimulates firing of dormant origins during recovery from HU ^15^, suppression of *de novo* origin firing with the CDC7 inhibitor XL413 (CDC7i) during HU did not reduce the increased association of POLA1 with ssDNA when RAD52 is inhibited as detected by PLA (Supplementary Figure 6). These findings argue against a possibility that Polα recruitment stimulated by loss of RAD52 depends on *de novo* origin firing in HU. After demonstrating that replicative gaps seen following loss of RAD52 are independent of PrimPol and that RAD52 inhibition stimulates Polα recruitment over PrimPol, we next examined whether inhibiting Polα could reverse the increase in parental ssDNA associated with the inhibition of RAD52i.

Inhibition of Polα dramatically increases ssDNA at fork and eventually blocks replication ^34^. Under HU conditions, where the majority of replication forks are delayed, even a low dose of inhibitor might affect the small amount of Polα involved in response to RAD52i. Thus, we decided to titrate the amount of a Polα inhibitor (ST1926; Polαi) against EdU incorporation and select a concentration that does not impact significantly on DNA synthesis. While the doses of the inhibitor previously used to block Polα ^34^ suppressed EdU incorporation, the 0.3µM dose only minimally reduced the number of positive cells and EdU immunofluorescence intensity in each nucleus (Supplementary Figure 7). Then, we evaluated the presence of parental ssDNA at 0.5mM HU in the presence of Polαi during treatment (Figure 2D). To exclude the contribution of origin-dependent function of Polα, we combined the low-dose of Polαi with the CDC7i. Wild-type cells were also treated with the higher dose of Polαi (3µM) as a positive control for ssDNA accumulation.

Consistent with an involvement of Polα in the accumulation of parental ssDNA when RAD52 is inhibited, treatment with a low dose of Polαi greatly decreased the amount of parental ssDNA in cells with inhibited RAD52, while not reducing ssDNA exposure in wild-type cells (Figure 2D). Combined treatment with the CDC7i and 0.3µM Polαi did no further reduce parental ssDNA during HU treatment of RAD52-inhibited cells or in control cells (Figure 2D), excluding the involvement of new origin firing. As expected, in unperturbed cells, the high dose of Polαi (3µM) led to the exposure of one order of magnitude more parental ssDNA if compared to the low dose (Figure 2D). These results indicate that the treatment with a low-dose of Polαi during perturbed replication can be used to target *de novo*, and origin-independent, recruitment of Polα in RAD52-inhibited cells. We next evaluated the formation of DNA gaps using a modified version of the alkaline Comet assay that is coupled with BrdU detection in nascent DNA ^35,36^. The BrdU alkaline Comet assay is able to detect ssDNA gaps even when these gaps are present on just one of the two newly synthesized complementary DNA strands. Inhibition of RAD52 led to a significantly higher tail moment in BrdU-positive cells compared to wild-type cells, indicating the presence of more ssDNA gaps after 0.5mM HU treatment (Figure 2E). Of note, and consistent with the parental ssDNA, the BrdU-labelled tails DNA in RAD52-inhibited cells were completely eliminated by treatment with the low dose of the Polαi (Figure 2E).

To further correlate Polα to the formation of replicative gaps in cells inhibited of RAD52, we performed the S1 DNA fiber assay in wild-type or PrimPol KO MRC5SV40 cells, treated or not with the low-dose of Polαi and with 0.5mM HU. Our expectation was to find PrimPol-dependent gaps in RAD52-proficient cells and Polαi-sensitive gaps in cells inhibited of RAD52. The S1 nuclease treatment did not affect the IdU/CldU ratio in wild-type cells (Figure 2F). When we compared the IdU/CldU ratio from fibers with and without prior S1 nuclease treatment, we found no significant change in wild-type cells even when POLA1 was inhibited (Figure 2F). In PrimPol KO cells, inhibition of POLA1 reduced the IdU/CldU ratio, possibly indicating that gaps were introduced but independently on the PrimPol or Polα function (Figure 2F). Compared with RAD52-proficient wild-type cells, inhibition of RAD52 reduced the IdU/CldU ratio of the S1-treated DNA fibers and the tract length ratio was not recovered by PrimPol KO (Figure 2F). In contrast, the low-dose of Polαi increased significantly the IdU/CldU ratio in S1 DNA fibers from RAD52i-treated wild-type cells and, in a similar extent, also in the PrimPol KO (Figure 2F).

Collectively, these results indicate that inhibition of RAD52 stimulates an origin-independent recruitment of Polα that is linked to accumulation of replicative gaps through a Polα-dependent but PrimPol-independent mechanism.

### Recruitment of Polα at perturbed replication forks depends on fork reversal

Loss or inhibition of RAD52 stimulates SMARCAL1 loading to replication forks and elevated degradation of the RFs ^15,16^. We next investigated if stimulation of Polα recruitment could occur downstream these events. Thus, we analysed association of Polα with parental ssDNA by PLA in MRC5SV40 cells stably expressing a doxycycline-regulated shSMARCAL1 cassette ^37^. Concomitant depletion of SMARCAL1 and RAD52 inhibition greatly reduced the recruitment of Polα at parental ssDNA (Figure 3A). Of note, downregulation of SMARCAL1 *per se* decreased the number of PLA spots, although not significantly. Consistent with the PLA data, depletion of SMARCAL1 reduced the amount of POLA1 detected in chromatin fractions in cells treated with the RAD52i (Figure 3B).

**Figure 3.**
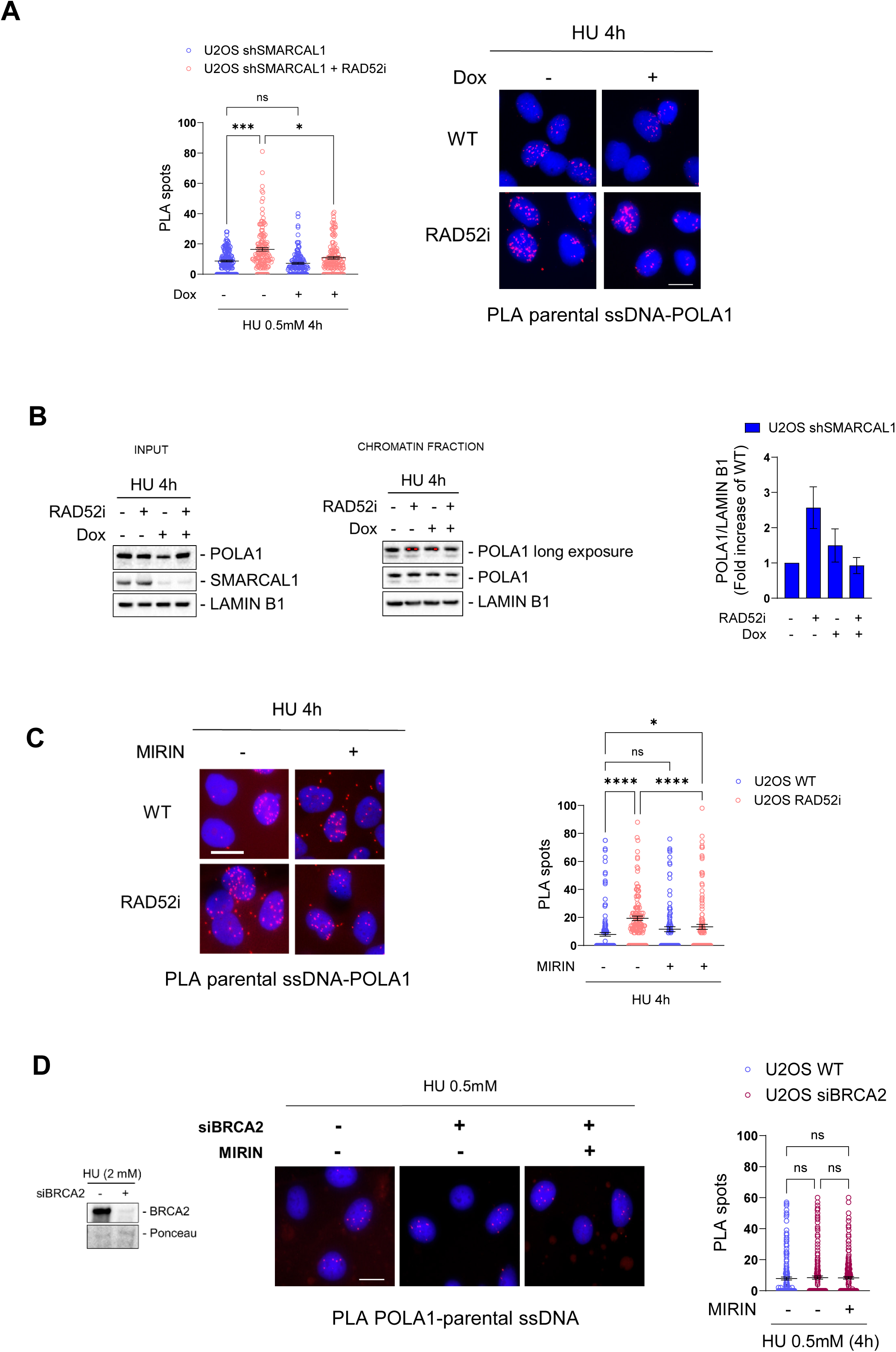
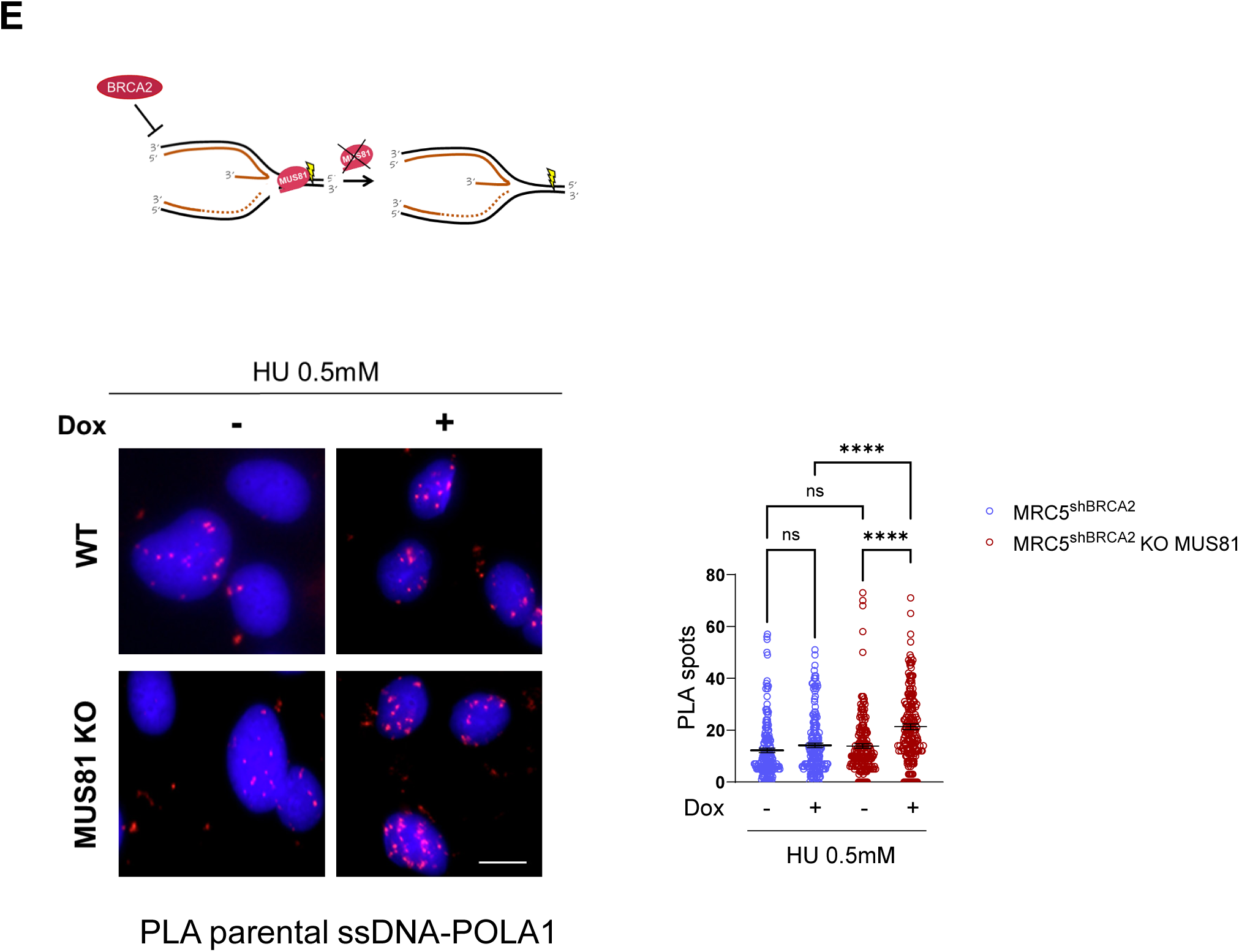
Polα recruitment under RAD52 deficiency depends on replication fork remodelling. **A.** Analysis of Polα-parental ssDNA interaction by anti-IdU/POLA1 PLA in inducible U2OS shSMARCAL1 cells. SMARCAL1 silencing was induced by giving Doxycycline (Dox) 24 hours before treatment with IdU for 20 hours. The cells were then released for 2 hours in fresh medium and treated with or without RAD52i 30 min before giving HU. Graph shows the number of PLA spot per nucleus. Representative images are shown. (ns = not significant; *P < 0.1; ***P < 0.001; Kruskal-Wallis test). Scale bar represents 10 µm. **B**. Analysis of Polα recruitment at DNA through chromatin fractionation in inducible U2OS shSMARCAL1 cells. Polα was identified by using an antibody directed against the Polα subunit POLA1, and LAMIN B1 used as loading control. Graph shows POLA1/LAMIN B1 quantification from 2 replicates. **C.** Analysis of the effect of MRE11 inhibition on Polα-parental ssDNA recruitment in RAD52i cells. PLA reaction was carried out using antibodies against POLA1 and IdU. The graph reports the number of PLA spot per nucleus. Representative images are shown. (ns = not significant; *P < 0.1; ****P < 0.0001; Kruskal-Wallis test). Scale bar represents 10µm. **D.** Analysis of Polα-parental ssDNA interaction in cells depleted of BRCA2. Western blot shows BRCA2 expression level after silencing with the siRNA. After IdU labelling, cells were treated with MIRIN and HU (0.5mM). PLA reaction was carried out using antibodies against POLA1 and IdU. Graph shows the number of PLA spots per nucleus. Representative images are shown. **E.** Analysis of Polα-parental ssDNA interaction in inducible shBRCA2 cells or in the MUS81 KO counterpart. BRCA2 silencing was obtained by treating the cells with Dox 48 hours before HU. PLA reaction was carried out using antibodies against POLA1 and IdU. Graph shows the number of PLA spots per nucleus. (ns = not significant; *P < 0.1; **P<0.1; ****P < 0.0001; Kruskal-Wallis test). Scale bar represents 10 µm.

We then investigated if the SMARCAL1-dependent recruitment of Polα stimulated by RAD52 inhibition was related to the nascent-strand degradation that occurs at RFs in this condition ^15,16^. Thus, we blocked fork degradation by exposing cells to the MRE11 inhibitor MIRIN, alone or in combination with the RAD52i, and then analysed the presence of POLA1 at parental ssDNA by PLA (Figure 3C). In cells where RAD52 was inhibited, the hyper-recruitment of POLA1 at ssDNA was significantly decreased by MIRIN but not completely eliminated. These results suggest that the hyper-recruitment of Polα occurring when RAD52 is inhibited is mostly a consequence of MRE11-dependent degradation at remodelled forks ^15^. To investigate if stimulation of Polα recruitment might represent a general response to the degradation of RFs, we analysed the presence of Polα at parental ssDNA by performing PLA experiments in BRCA2-depleted cells; the classical model of fork deprotection and MRE11-dependent degradation. BRCA2 depletion did not increase POLA1 association with parental ssDNA during replication fork arrest, nor was this association altered by interfering with MRE11-related fork degradation (Figure 3D).

Loss or inhibition of RAD52 leads to fork degradation but does not induce DNA breaks ^15,38^. This differs from the absence of BRCA2, where degraded RFs are subsequently converted into DSBs by the MUS81 complex ^39^. To test if cleavage of the degraded RFs by the MUS81 complex is the event preventing the recruitment of Polα, we performed parental ssDNA-POLA1 PLA in shBRCA2 cells KO for MUS81 ^40^. In these cells, MUS81-dependent DSBs at degraded forks are prevented as it occurs in the absence of RAD52 (scheme in Figure 3E). When BRCA2 was downregulated in cells with functional MUS81 (WT; + Dox), there was no change in the number of POLA1-ssDNA PLA spots. However, in shBRCA2/MUS81 KO cells, a clear increase in POLA1-ssDNA PLA spots was observed (Figure 3E).

Altogether, these results indicate that, in response to perturbed replication the absence of RAD52 stimulates *de novo* recruitment of Polα downstream fork reversal and degradation. They also suggest that this *de novo* recruitment of Polα downstream fork reversal and degradation correlate with failure to cleave the structure by the MUS81 complex.

### The function of Polα at perturbed replication forks involves interaction with RAD51

We observe that recruitment of Polα in response to loss of RAD52 depends on fork remodelling by SMARCAL1. In Xenopus extracts, Polα associates with Rad51 after replication arrest ^41^. Thus, we tested if origin-independent recruitment of Polα that occurs following inhibition of RAD52 might involve association with RAD51 as well. To this end, we first assessed the interaction between Polα (POLA1) and RAD51 by PLA at different time points after treatment with 0.5mM HU. Association of Polα with RAD51 was observed already in wild-type cells after replication perturbation, and increased with time becoming more evident at 4h of HU (Figure 4A). Interaction between Polα and RAD51 was higher and statistically significant in cells treated with the RAD52i at both 2 and 4h of HU (Figure 4A).

**Figure 4.**
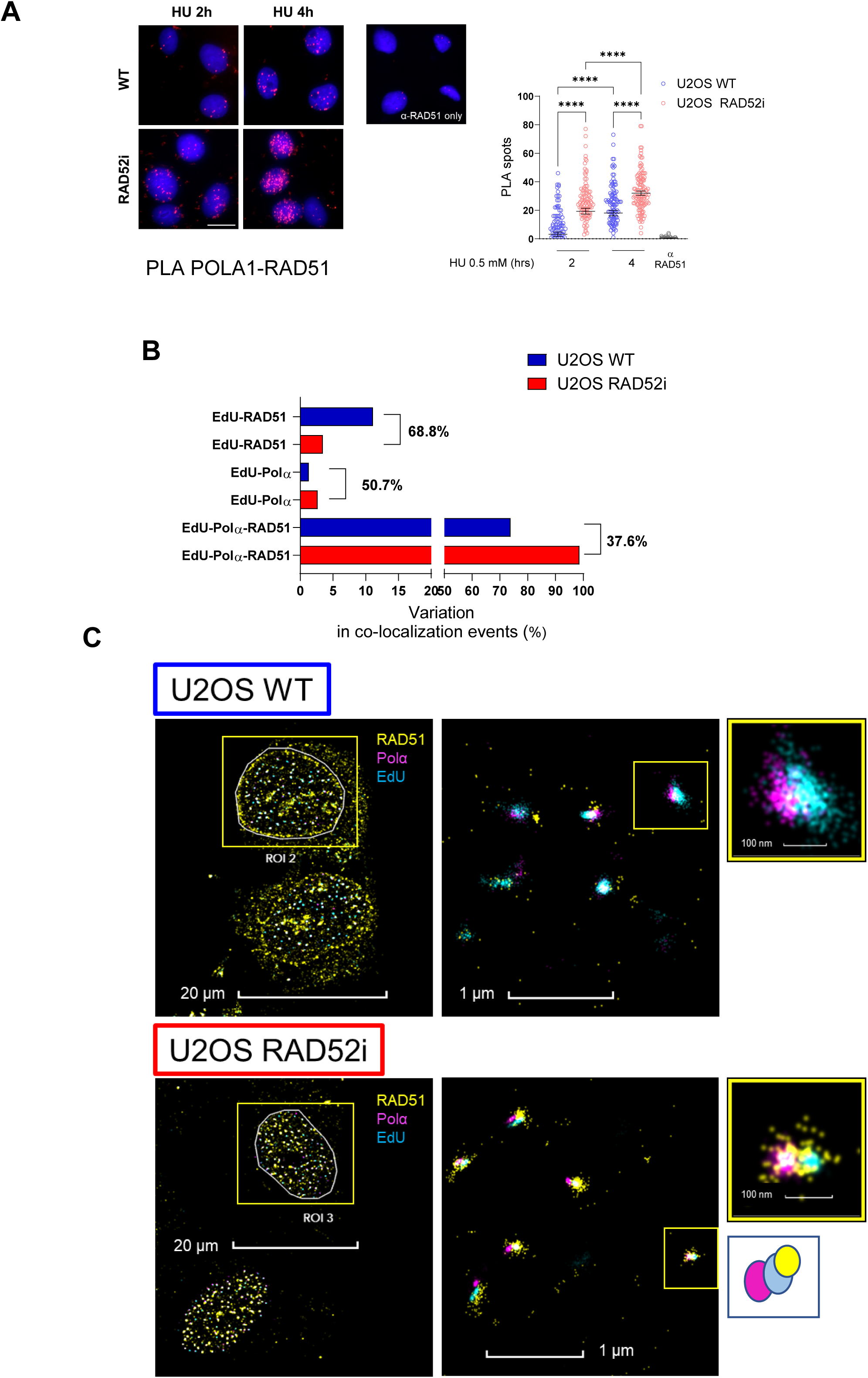
RAD52 inhibition increases Polα-RAD51 association. **A.** Analysis of Polα-RAD51 association by PLA in U2OS WT. Cells were subjected to RAD52i 30 min before HU treatment. PLA reaction was carried out using antibodies against POLA1 and RAD51. The graph shows the number of PLA spot per nucleus. The control was subjected to PLA with only one primary antibody. Representative images are shown. Scale bar represents 10 µm. **B.** Quantification of EdU-RAD51-Polα interaction by dSTORM nanoscopy in U2OS cells. The graph represents the percentage of variation in co-localization events of EdU-RAD51, EdU- Polα and EdU-RAD51-Polα in localization clusters. **C.** Representative dSTORM images of two nuclei immunolabeled for nascent DNA (cyan), Polα (magenta) and RAD51 (yellow). Scale bars = 20µm, 1µm and 100nm. A representative cartoon of a common topology of the 3 signal is also shown. All the values above are presented as means ± SE (ns = not significant; *P < 0.1; **P<0.1; ***P < 0.001; ****P < 0.0001; Kruskal-Wallis test).

To further confirm that RAD52 inhibition stimulated Polα recruitment and association with RAD51 at perturbed replication forks, we performed correlative single-molecule localisation microscopy by dSTORM. Active replication forks were labelled with a short EdU pulse before challenging replication with 0.5mM HU in the presence or not of RAD52i (Figure 4B). After filtering out single labelling sites, co-localisation at the nanometry-scale was assessed in tricolour dSTORM through the analysis of the variation of signal clustering involving EdU-Polα or Polα-RAD51 at EdU+ sites (Figure 4B). These analyses confirmed the increase in the fraction of Polα at EdU+ sites observed with PLA and revealed that RAD52 inhibition resulted in a shift of the fraction of RAD51 molecules that engage in single interaction with the fork to the benefit of the fraction involved in the association with Polα at the EdU+ sites (Figure 4B, C). Furthermore, topology inspection suggested that Polα is found almost always at the end of the EdU signal that is preceded by or embedded into the RAD51 signal in the clustered analysis. Since the number of EdU clusters appeared higher than that of Polα-RAD51-EdU clusters, this suggests that not all perturbed replication forks utilize this repriming pathway when RAD52 is inhibited, indicating more than a single alternative mechanism involved.

Since RAD52 inhibition led to increased Polα recruitment, we sought to determine whether this effect depended on RAD51 activity. To address this, we evaluated the association of Polα with parental ssDNA by PLA in cells treated or not with the RAD51 inhibitor B02 (RAD51i; ^42^). As shown in Figure 5A, RAD51 inhibition increased the association of Polα with parental ssDNA at 2h of HU treatment in RAD52-proficient cells (WT). However, this effect was not significant at 4h of treatment. In contrast, RAD51 inhibition led to a striking reduction in the number of PLA spots when RAD52 was inhibited. Notably, concomitant inhibition of RAD51 and RAD52 returned the level of Polα-ssDNA interaction to that detected in wild-type cells (Figure 5A). Most importantly, concomitant inhibition of RAD51 and RAD52 greatly reduced the increased formation of parental ssDNA (Supplementary Figure 8).

**Figure 5.**
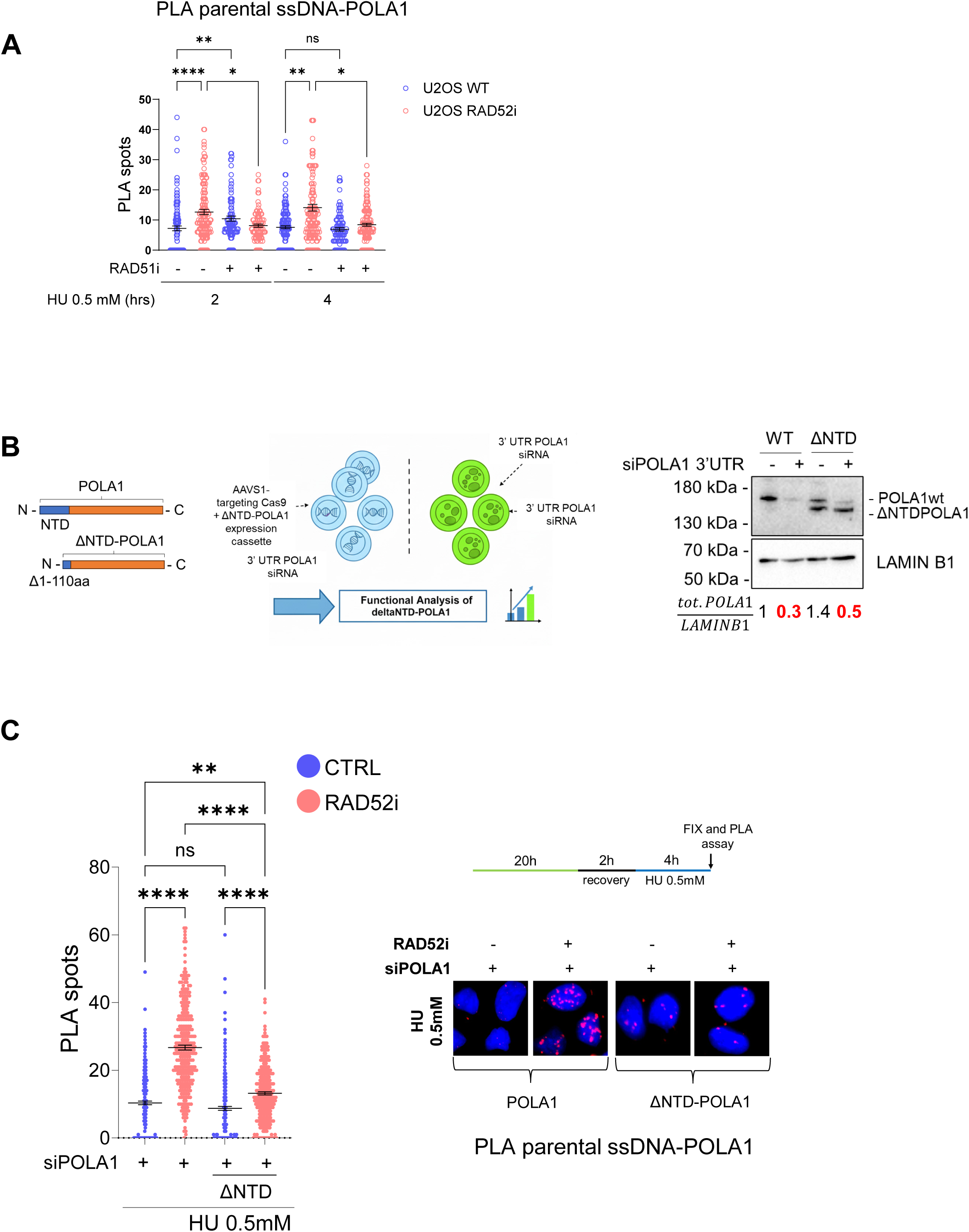
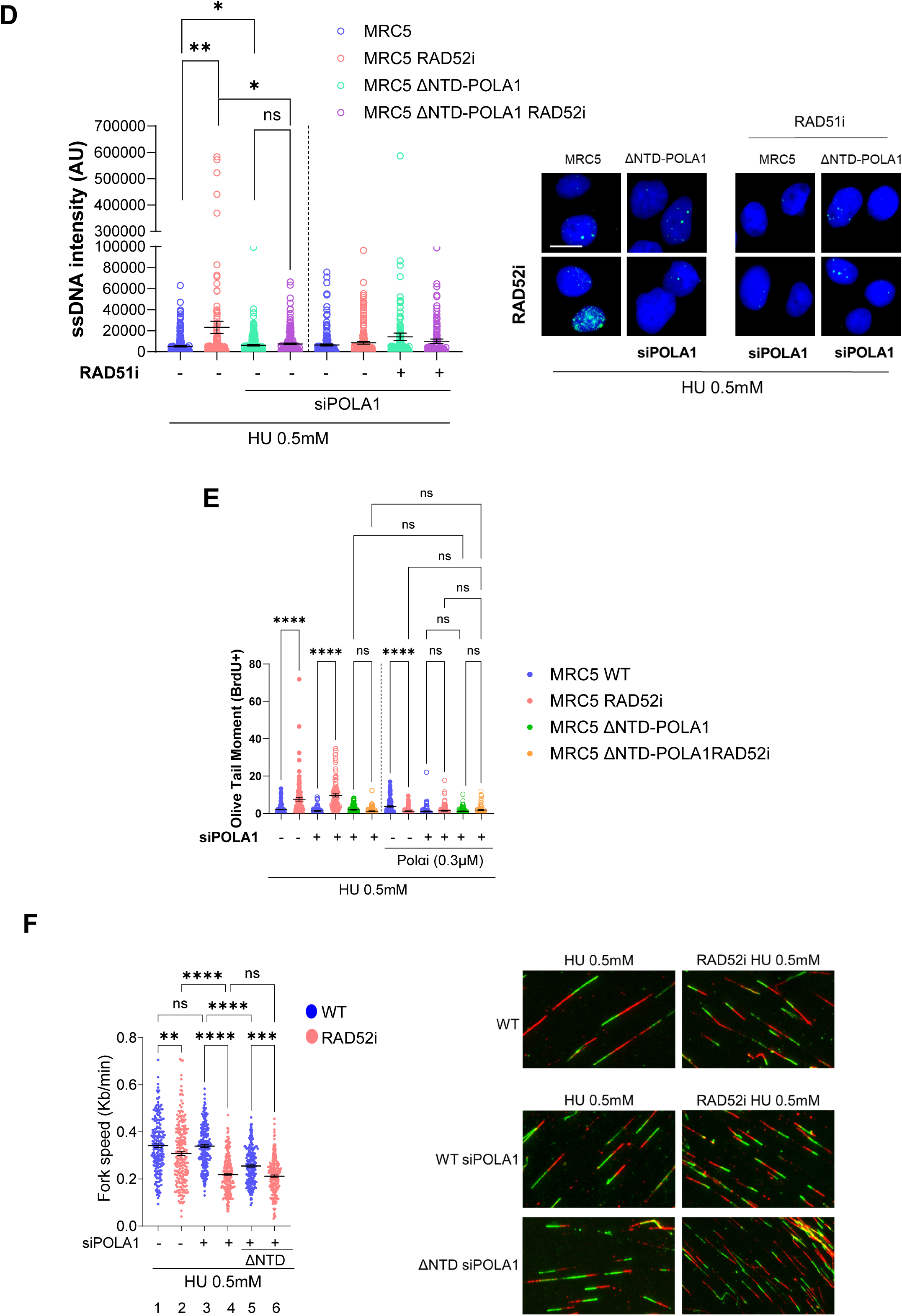
Polα engagement is mediated by RAD51. **A.** Analysis of Polα-parental ssDNA association by PLA. U2OS cells were treated with IdU for 20 hours, released for 2 hours in fresh DMEM and successively treated with or without RAD52i and RAD51i 30 min before HU. The graph shows the number of PLA spot per nucleus. All the values above include means ± SE (ns = not significant; *P < 0.1; **P<0.1; ****P < 0.0001; Kruskal-Wallis test). **B**. Cartoon representing the generation of the ΔNTD-POLA1-expressing cells. ΔNTD-POLA1-expressing MRC5 cells were generated by gene-targeting mediated by Cas9 at the AAVS1 site and selected to identify homozygous editing. For functional analysis, cells were transfected with siRNA directed against the 3’ UTR of POLA1 as indicated in the scheme. Western blot analysis shows level of protein after 48h of transfection with control or POLA1 siRNA. The total POLA1 ratio represents the total amount of POLA1 (endogenous + ΔNTD). In red the level of POLA1 in the cell backgrounds used in functional experiments. **C.** Analysis of Polα-parental ssDNA association by PLA in ΔNTD-POLA1-expressing cells. The graph shows the number of PLA spot per nucleus. Representative images are reported. (ns = not significant; ***P < 0.001; ****P < 0.0001; Kruskal–Wallis test). Scale bar represents 10 µm. **D.** Analysis of parental ssDNA exposure by immunofluorescence. MRC5 WT and MRC5 ΔNTD-POLA1 cells were transfected with POLA1 siRNA as indicated in **B**. After the exposure to HU, the ssDNA was detected by native anti-IdU immunofluorescence. Graph shows the intensity of ssDNA staining (AU) per cell. Representative images are shown. (ns = not significant; *P < 0.1; **P<0.1; Kruskal–Wallis test). Scale bar represents 10 µm. **E.** BrdU alkaline Comet assay. MRC5 WT and MRC5 ΔNTD-POLA1 cells were transfected with POLA1 siRNA as indicated in **B**. Polαi was added 30 min before HU (0.5mM). Graph shows the individual tail moments in BrdU+ cells from at least 50 nuclei from 3 independent repeats. (ns = not significant; *P < 0.1; **P<0.1; Kruskal–Wallis test). **F.** Analysis of fork progression by DNA fiber assay in MRC5 WT and MRC5 ΔNTD-POLA1 cells transfected with POLA1 siRNA as indicated in **B**. The graph shows the individual fork speed values from 3 independent repeats. Representative images are shown. (ns = not significant; **P<0.1; ***P < 0.001; ****P < 0.0001; Kruskal-Wallis test).

Experiments with Xenopus egg extracts indicate that RAD51 interacts with the NTD region of POLA1 ^41^. Thus, we generated MRC5SV40 cells expressing a truncated version of POLA1 lacking the first 110aa of the protein containing the RAD51-binding region (ΔNTD-POLA1) and silenced endogenous POLA1 using RNAi (Figure 5B). As shown in the WB of Figure 5B, the ΔNTD-POLA1 was not overexpressed and RNAi reduced the amount of endogenous wild-type POLA1 to that of the ectopic ΔNTD-POLA1. Both wild-type cells expressing reduced levels of endogenous POLA1 and those expressing ΔNTD-POLA1 were viable and showed only minor reduction in the proliferation rate compared to the mock-silenced (Supplementary Figure 9A). Next, we investigated if ΔNTD-POLA1 was really unable to associate with RAD51 when RAD52 was inhibited. As expected, RAD51-POLA1 PLA confirmed that RAD52-inhibited cells expressing the ΔNTD-POLA1 and silenced for endogenous, wild-type, POLA1 showed minimal interaction between POLA1 and RAD51 after replication stress (Supplementary Figure 9B). Using this experimental model, we tested the association of Polα with parental ssDNA by PLA after treatment with 0.5mM HU, and in the presence or not of the RAD52i. Although the level of POLA1 was reduced by RNAi, RAD52 inhibition still stimulated Polα recruitment at parental ssDNA (Figure 5C). Most notably, expression of ΔNTD-POLA1 greatly reduced recruitment of Polα in RAD52-inhibited cells but not in the control cells, as shown by the decrease in the number of PLA spots (Figure 5C). Since the inhibition of RAD51 is sufficient to prevent accumulation of parental ssDNA in RAD52-deficient cells, we evaluated if expression of the ΔNTD-POLA1, which interferes with interaction with RAD51, could prevent this phenotype and mimicking the effect of a low-dose of Polα inhibitor (see Figure 2D and E). RAD52 inhibition led to accumulation of parental ssDNA also in wild-type cells expressing a reduced amount of POLA1 because of the RNAi (Figure 5D). Expression of ΔNTD-POLA1, largely prevented accumulation of parental ssDNA in RAD52-inhibited cells, mimicking the effect of the RAD51i (Figure 5D). Consistent with the parental ssDNA analysis, the BrdU alkaline Comet assay showed that expression of the ΔNTD-POLA1 protein suppressed accumulation of replication stress-associated DNA gaps in the daughter strand when RAD52 is inhibited (Figure 5E). Interestingly, treatment with the low dose of Polαi (0.3µM) did not further affect the detection of DNA gaps in RAD52-inhibited cells expressing the ΔNTD-POLA1 protein (Figure 5E).

Inhibition of RAD52 did not affect fork rates in untreated cells or recovery from 2mM HU (Supplementary Figure 3 and ref. ^15^). Thus, we tested if this RAD51 and Polα-mediated pathway could be required for fork progression under perturbed replication when RAD52 was inhibited. To this aim, we performed DNA fibre assay in the cell model expressing the ΔNTD-POLA1 protein that cannot interact with RAD51 and consequently does not support repriming. As compared with non-inhibited cells, shown in Figure 5F, inhibition of RAD52 inhibition reduced fork progression in cells treated with 0.5mM HU even if they expressed the endogenous POLA1 (Figure 5F, 2 vs 1). When RAD52 is functional, even expression of a reduced amount of POLA1 did not affect fork progression under perturbed conditions (Figure 5F, 1 vs 3). In contrast, even expression of a reduced amount of endogenous, wild-type, POLA1 was sufficient to affect fork progression when RAD52 was inhibited (Figure 5F, 2 vs 4). Similarly, concomitant RAD52 inhibition and expression of ΔNTD-POLA1 led to a strong impairment of fork progression that was comparable to the effect of the reduced amount of POLA1 (Figure 5F, 2 vs 6 vs 4). Unexpectedly, expression of ΔNTD-POLA1 reduced fork progression significantly under perturbed replication even when RAD52 was functional (Figure 5F, 3 vs 5), possibly suggesting that the interaction between RAD51 and Polα is required also independently of loss of RAD52-mediated fork protection.

These results indicate that the origin-independent Polα recruitment and parental ssDNA accumulation stimulated by loss of RAD52 function during replication perturbation are dependent on the interaction of POLA1 with RAD51. Furthermore, our results using the ΔNTD-POLA1 cells indicate that the RAD51-dependent Polα repriming is required for fork recovery when RAD52 is inhibited.

### Polα recruitment occurs downstream extended degradation at perturbed replication forks and requires stable RAD51 nucleofilaments

Our data suggest that the engagement of Polα when RAD52 is inhibited involves RAD51 nucleofilaments assembly. To test the hypothesis that RAD51-mediated Polα recruitment during perturbed replication requires extensive nucleolytic degradation of the RF, we performed pulse and chase SIRF experiments. This experiment aimed to define whether Polα can be found close to EdU-labelled nascent DNA following degradation of the unlabelled nascent DNA at RF or behind it (see scheme in Figure 6A). Thus, we pulse-labelled active replication forks with EdU followed by extensive washing and chase in EdU-free medium supplemented with Thymidine and 0.5mM HU, to slow-down replication, in the presence or absence of MIRIN and RAD52i (Figure 6A). Assuming 1/2 to 1/10 of normal replication fork rate in 0.5mM HU (∼ 0.1-5 vs. 1Kb/min; ^43^), these pulse and chase experiments were expected to locate labelled nascent DNA far away to the RF (Figure 6A). At the 15min chase time-point, more Polα became associated with EdU-labelled DNA in RAD52 inhibited cells. However, this increased association was completely suppressed by MIRIN in addition to RAD51i, which set the PLA signals back to the wild-type values (Figure 6B).

**Figure 6.**
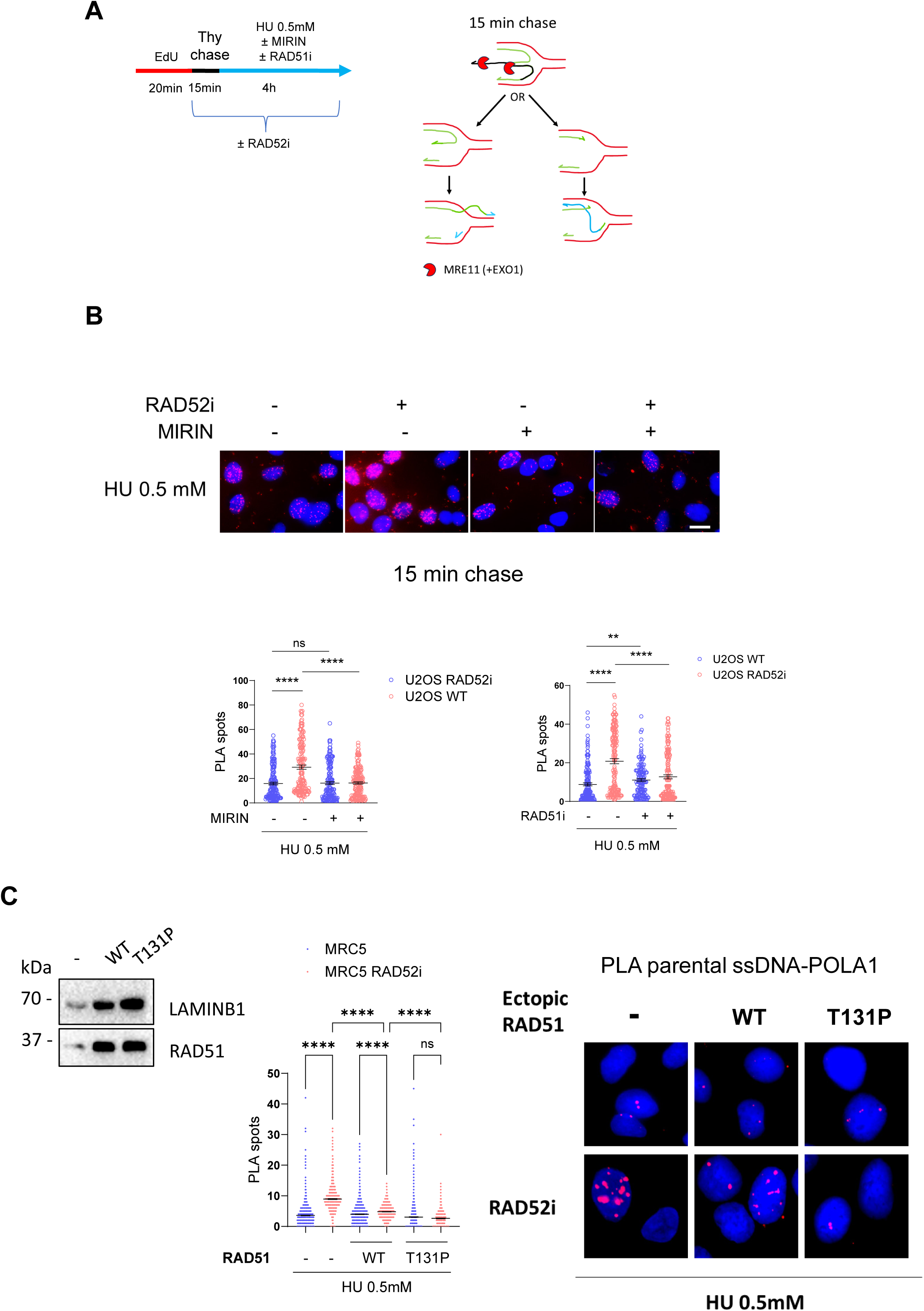
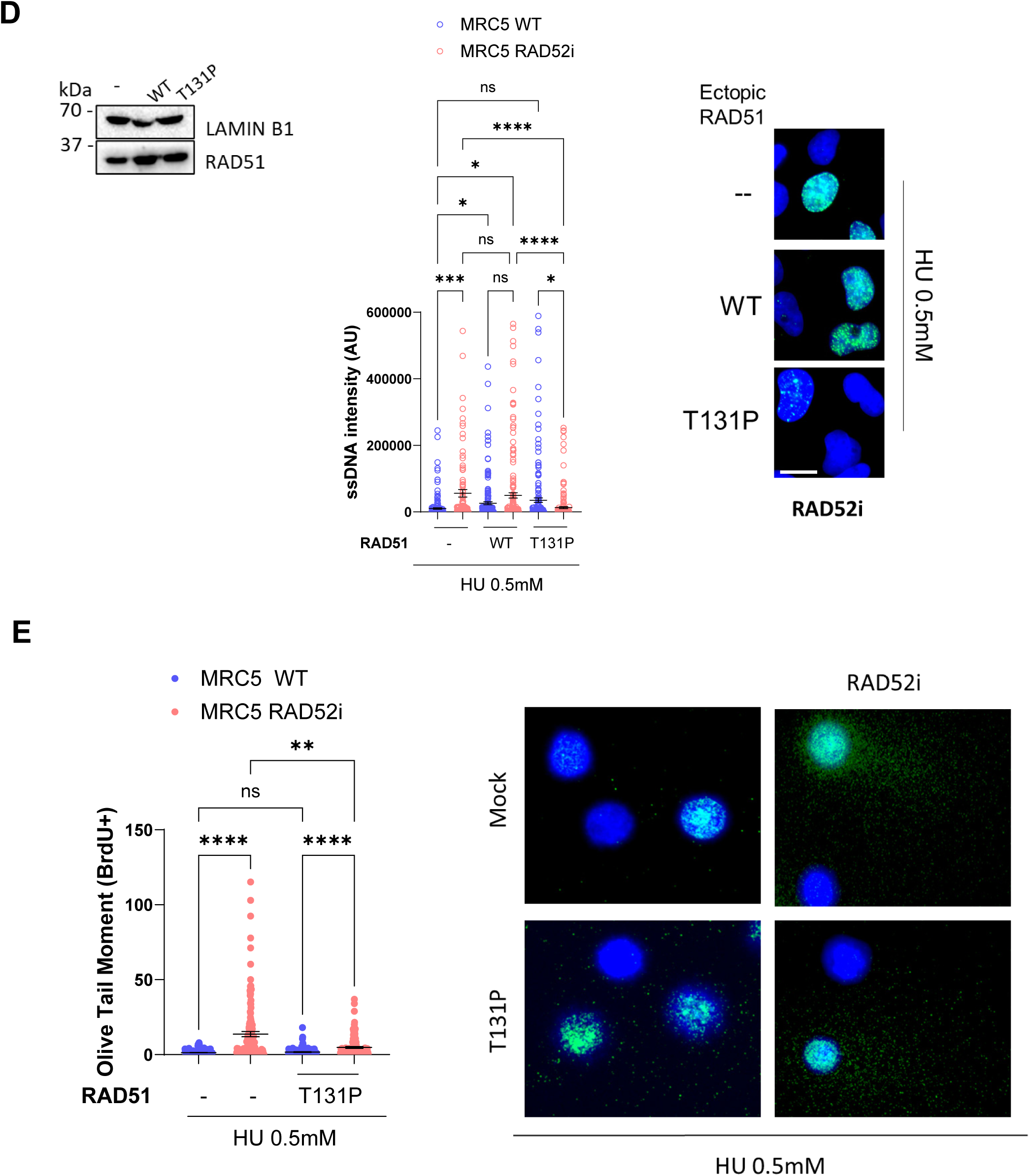
Polα recruitment occurs downstream extensive replication fork degradation and requires RAD51 nucleoprotein filaments. **A.** Experimental scheme of the pulse/chase analysis of Polα-nascent ssDNA interaction by SIRF assay in U2OS cells. The cartoon depicts the expected localisation after the chase. **B.** The graph shows the number of EdU-POLA1 spots per nucleus from 3 different repeats. (ns = not significant; **P<0.1; ****P < 0.0001; Kruskal-Wallis test). The representative images are shown above the graph. Controls were subjected to SIRF with anti-biotin only. **C.** QIBC analysis of Polα-parental ssDNA interaction in cells expressing the RAD51 T131P mutant. Cells were transfected with RAD51 WT or T131P and parental DNA labelled with IdU 24h after transfection. After 20h cells were released in fresh medium for 2h and treated with HU for 4 hrs. Western blot shows RAD51 WT and RAD51 T131P in transfected cells. LAMIN B1 was used as a loading control. Graph shows the number of PLA spots from 3 replicates. (ns = not significant; ****P < 0.0001; Kruskal-Wallis test). **D.** Analysis of parental ssDNA exposure in MRC5 WT cells transfected or not with RAD51 WT or RAD51 T131P. Western blot shows level or RAD51 in transfected cells. Graph shows the intensity of ssDNA staining (AU). (ns = not significant; *P<0.5; ***P < 0.001; Kruskal-Wallis test). Representative images are shown. **E.** BrdU alkaline Comet assay in cells with impaired formation of RAD51 nucleoprotein filaments. Cells are treated and processed as explained in the Figure 2E. The graph shows the individual tail moments in BrdU+ cells from 3 repeats. (ns = not significant; **P<0.1; ****P < 0.0001; Kruskal–Wallis test). Representative images are shown.

These results suggest that defective metabolism of stalled forks in RAD52-inhibited cells promotes a RAD51-dependent recruitment of Polα that is downstream of extensive fork degradation, and that might implicate strand invasion from the gap formed at the reset RF to promote recombination-dependent replication. As such, recruitment of Polα should occur downstream RAD51 nucleofilament and D-loop formation. To test this hypothesis, we expressed the wild-type or the T131P mutant of RAD51 by transient transfection in MRC5SV40 cells and analysed the recruitment of Polα at parental ssDNA by POLA1-IdU PLA after treatment with 0.5mM HU (Figure 6C). The RAD51-T131P mutant, when expressed together with the endogenous wild-type protein, forms unstable nucleofilaments but no D-loops and it is also known to induce fork destabilisation ^44,45^. However, we know that fork destabilisation *per se* is not sufficient to stimulate recruitment of Polα (see Figure 3D). Therefore, in the presence of RAD52i, the expression of RAD51-T131P should efficiently test if stable nucleofilaments and D-loop formation are essential for the subsequent recruitment of Polα. Recruitment of Polα at parental ssDNA by PLA was quantified using QIBC in S-phase-gated populations. As shown in Figure 6C, inhibition of RAD52 recapitulated the increased interaction of Polα with parental ssDNA. Overexpression of wild-type RAD51 apparently did change the recruitment of Polα in the absence of RAD52 inhibition while preventing recruitment of Polα in cells inhibited of RAD52 (Figure 6C), consistent with the reported rescue of the fork degradation phenotype^15^. In RAD52-inhibited cells, the expression of RAD51-T131P suppressed the recruitment of Polα and restored the same level of PLA spots seen in mock-transfected wild-type cells (Figure 6C). Consistent with this effect, the expression of RAD51-T131P also prevented the association between RAD51 and Polα in RAD52-inhibited cells (Supplementary Figure 10).

We next investigated if expression of RAD51-T131P was sufficient to reduce parental ssDNA accumulation and DNA gap formation in RAD52-inhibited cells. Thus, we performed native IdU detection after parental labelling in cells treated with 0.5mM HU in the presence of the RAD52i and the ectopic wild-type or T131P form of RAD51 (Figure 6D). As expected, inhibition of RAD52 increased the exposure of ssDNA in the parental strand, a sign of daughter-strand gaps (Figure 6D). The presence of more wild-type RAD51 in the mock-inhibited cells stimulated the exposure of parental ssDNA, consistent with more Polα detected at parental ssDNA by PLA, while it did not affect exposure of parental ssDNA in RAD52-inhibited cells (Figure 6D). However, and consistent with the reduction of Polα recruitment, ectopic expression of the RAD51-T131P mutant prevented the RAD52i-related accumulation of parental ssDNA (Figure 6D).

To further substantiate our observations, we exposed mock or RAD52i-treated cells to 0.5mM HU for 1h and performed BrdU Comet assay to detect DNA gaps in the presence of T139P RAD51. Consistent with the parental ssDNA, overexpression of T131P RAD51 in mock-inhibited cells did not affect the very low Comet tail moment values in BrdU-positive cells implying no DNA gap formation (Figure 6E). In RAD52-inhibited cells, RAD51-T131P overexpression decreased the Comet tail moment in BrdU-positive cells, nearly eliminating DNA gap accumulation (Figure 6E).

Collectively, these results indicate that the Polα recruitment and DNA gap formation stimulated by loss of RAD52 activity occurs after extensive nucleolytic degradation of the RF and requires formation of stable RAD51 nucleofilaments.

### RAD51 nucleofilament formation directly stimulates Polα/Primase function

Our data indicate that RAD52 inhibition promotes RAD51-mediated Polα recruitment following replication perturbation, leading to the formation of daughter-strand gaps. Given that RAD51 and Polα interact in the cell, and this interaction is required for Polα-recruitment at distressed forks (Figure 5), we investigated whether RAD51 stimulates Polα-mediated repriming by testing its primase-polymerase activities through in vitro assays. To this end, we conducted a primer extension experiment using a 70nt-long model poly-dT template and recombinant Polα-Primase complex, with and without increasing amounts of RAD51. Reaction conditions were optimized to assess only the primase reaction while suppressing DNA synthesis (Supplementary Figure 11A-D). The purified Polα/Primase complex successfully performed priming, resulting in the expected size of primed DNA (Figure 7A). Adding RAD51 to the reaction surprisingly increased the efficiency of the primase function, extending up to the full length of the template and, at higher concentrations of RAD51, even beyond (Figure 7A-C), suggesting the joining of multiple templates by RAD51 nucleofilaments. RAD51 is a single-strand DNA binding protein. Thus, to determine if the observed stimulation of priming activity of Polα/Primase by RAD51 could be dependent on its ssDNA binding activity, we repeated the primer extension assay with purified RPA. As shown in Figure 7D-F, RPA completely suppressed the primase activity. Similarly, inclusion of RAD51 at increasing concentrations stimulated primase activity of Polα/Primase and formation of extra-long RNA products also using a more complex template DNA, the φX174 virion ssDNA, which is more than 5kb-long and circular (Supplementary Figure 12A-C). The stimulation by RAD51 was limited to the primase activity since switching into DNA synthesis-permissive conditions failed to show any stimulation of Polα-dependent DNA synthesis, which was reduced in the presence of RAD51 or RPA (Supplementary Figure 12D-F). We next tested the effect of RAD51 on the ability of Polα to perform primer extension at a D-loop (Figure 7G). We observed that Polα alone barely shows primer extension ability on the D-loop substrate while readily extends primer from a linear template. Notably, RAD51 facilitated primer extension by Polα on the D-loop structure (Figure 7H, K). Despite this stimulation being limited to no more than 20%, it appeared to be dependent on the concentration of RAD51 (Figure 7K). In contrast, the presence of RAD51 counteracted the primer extension activity of Polα on the linear template (Figure 7j, K).

**Figure 7.**
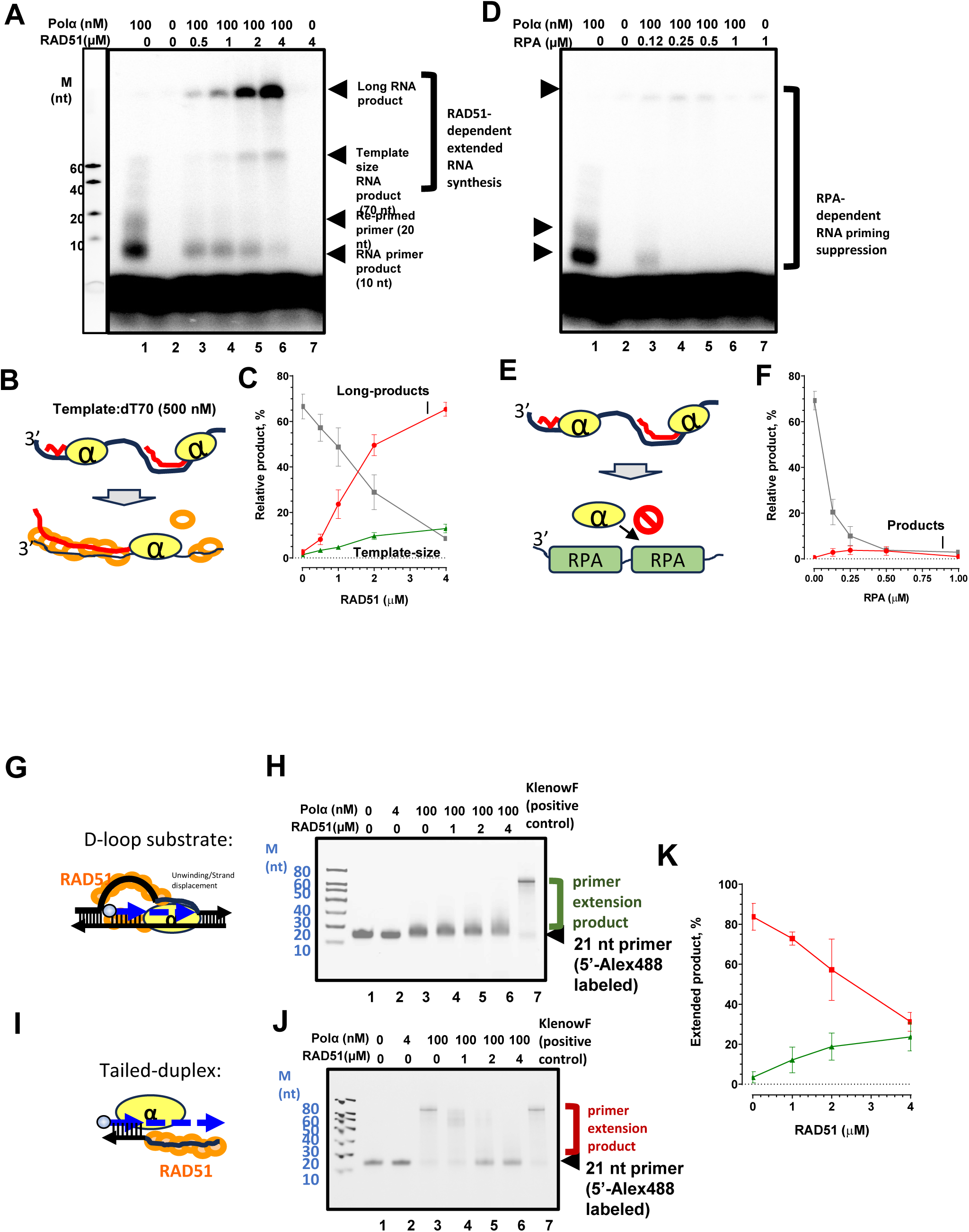
RPA and RAD51 differentially regulate Polα/Primase activities. **A.** RAD51 titration on Polα/Primase *de novo* RNA synthesis on dT70 mer templates, with products labelled with [α-^32^P]ATP. Size markers are synthesized Cy3-labelled poly(dT) ssDNA with a length of 10, 20, 40 and 60. **B.** Model depicting the observed elongated RNA products in the presence of RAD51 (orange circle). Polα–primase (yellow circle labeled as α) binds the template poly(dT)70 and synthesize RNA primer (red) that length could be modulated by RAD51. **C.** Quantification of RNA products in the experiment directly above. 10 and 20 nt RNA priming product (grey), template size RNA product (green) and long RNA product that is stacked in the well (red). **D.** RPA titration on Polα/Primase *de novo* RNA synthesis on dT70 mer templates, with products labelled with [α-^32^P]ATP. **E.** Model depicting the observed elongated RNA products in the presence of RPA (green square). Polα–primase (yellow circle labeled as α) binds the template poly(dT)70 and synthesize RNA primer (red) while its activity is attenuated in the presence of RPA. **F.** Quantification of RNA products in the experiment directly above. 10 and 20 nt RNA priming product (grey) and long RNA product (red). **G.** Model illustrating DNA primer extension on a D-loop structure in the presence of RAD51 (orange circle). The activity of Polα–primase (yellow circle labeled “α”) may be modulated by RAD51 either directly through physical interaction or indirectly via RAD51-mediated strand invasion and dsDNA unwinding, facilitating primer extension. **H.** A 5′-Alexa488-labeled 21-nt primer was extended by Polα–primase on a D-loop-mimicking substrate (green) in the presence of dNTP and varying amounts of RAD51. Primer extension can be carried out efficiently by the Klenow Fragment alone. **I.** Model showing DNA primer extension on a tailed-duplex substrate. Polα–primase efficiently extends the primer under this condition, but its activity is attenuated in the presence of RAD51. **J.** Same experimental setup as in H, but using a tailed-duplex substrate. Extension products are indicated in red. **K.** Quantification of DNA primer extension products from experiments b and d. Products from the D-loop substrate are shown in green; those from the tailed-duplex are shown in red.

Our cell-based experiments showed that the T131P RAD51 mutation inhibits Polα recruitment and parental ssDNA accumulation after RAD52 inhibition (Figure 6C-E). Therefore, we tested if proper RAD51 nucleofilament formation was needed for the observed increase in Polα/Primase activity using the poly(dT) template. To this end, we used RAD51-T131P and one additional mutant that is unable to assembly nucleofilaments (RAD51-F86E) ^46,47^. As shown in Supplementary Figure 13A and B, both RAD51 mutants abrogated, partially or completely depending on the level of impaired nucleofilament formation, the effect on the Polα/Primase activity on template ssDNA (Supplementary Figure 13C).

These findings show that RAD51 nucleofilaments directly enhance the primase activity of Polα, promoting primer extension at D-loops and enabling Polα/Primase to synthesize unusually long RNA primers.

### RAD51/Polα-dependent repriming is involved in limiting the accumulation of chromosomal damage and cell death in RAD52-deficient cells

We demonstrated that inhibition of RAD52 during replication fork perturbation stimulates a peculiar, origin-independent but RAD51-dependent, engagement of Polα that ensures fork progression under perturbed replication. Thus, we asked whether, despite generating DNA gaps, this mechanism might be protective against DNA damage and genome instability accumulation.

To this aim, we performed γH2AX immunostaining, a surrogate marker for DSBs, after replication perturbation in mock (WT) or RAD52-inhibited U2OS cells. To interfere with the Polα-mediated repriming events, we exposed cells to the low dose of POLA1i during treatment (see Figures 2D-F). As shown in Figure 8A, neither RAD52i nor POLA1i alone significantly increased the yield of γH2AX-positive cells in response to 0.5mM HU. However, combined with the RAD52i, treatment with POLA1i greatly enhanced the number of γH2AX-positive cells and the intensity of the staining.

**Figure 8.**
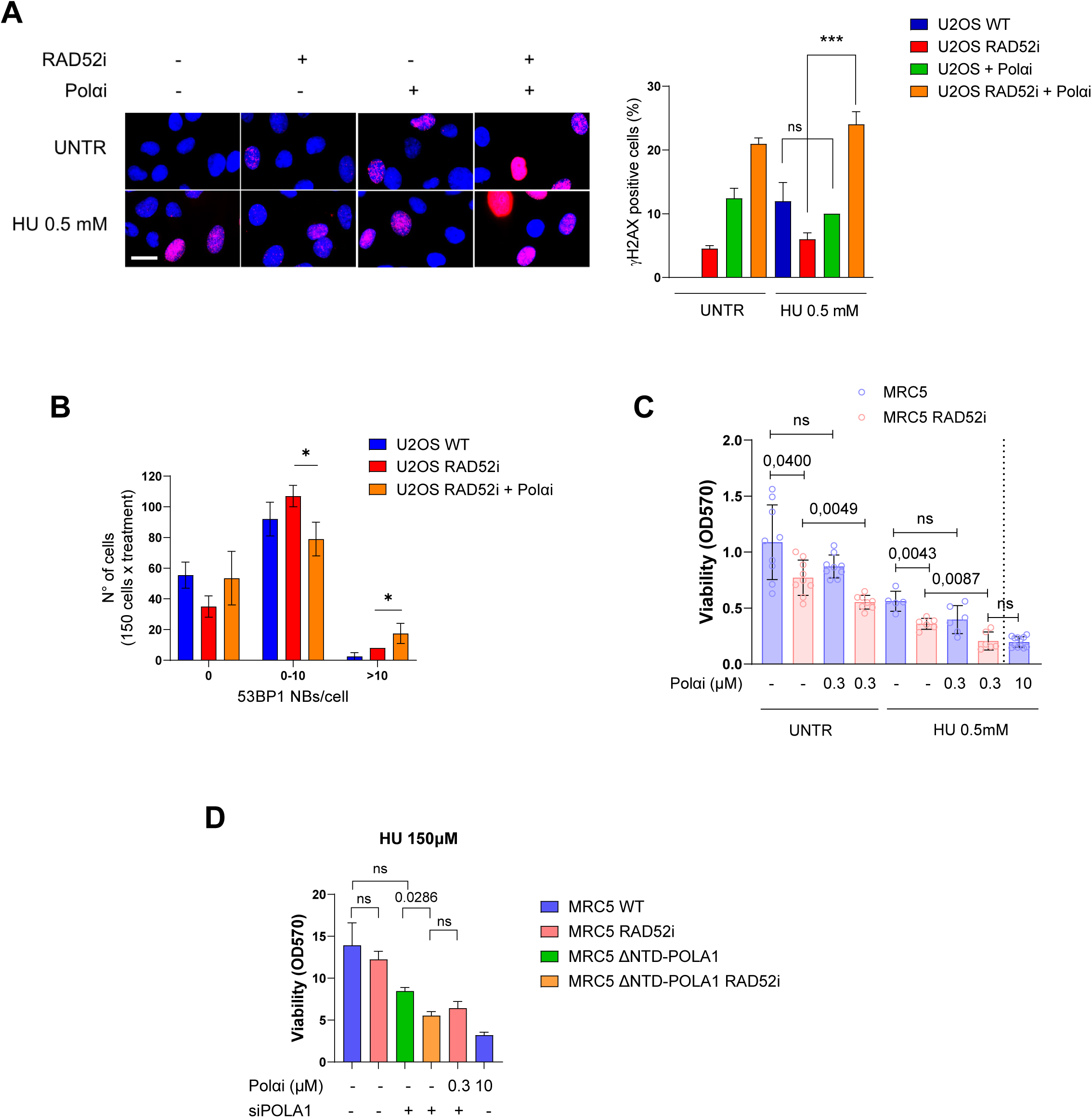
Polα-repriming prevents DSBs and chromosomal aberrations, ensuring viability in RAD52 deficient cells. **A.** DSBs detection by immunofluorescence. U2OS WT were treated for 4h with HU and inhibitors, and recovered for 18h in drug-free medium before analysis. Immunofluorescence was carried out by using an antibody against γH2AX. The graph shows the percentage of γH2AX positive cells. Representative images are shown. **B.** 53BP1 NBs detection. Cells were treated as indicated in the experimental scheme above. Immunofluorescence was carried out by using an antibody against 53BP1. The graph shows the number of cells presenting 53BP1 NBs (0, 0-10 or >10). All the values above are presented as means ± SE (ns = not significant; *P < 0.1; **P<0.1; ***P < 0.001; ****P < 0.0001; ANOVA test;). Scale bars represent 10 µm. **C-D.** Viability assay to detect effects of impaired RAD51-Polα pathway in RAD52-inhibited cells upon replication fork perturbation. Inhibitors and HU were maintained in the culture medium until fixation. All the values above are presented as means ± SE (ns = not significant; *P < 0.1; **P<0.1; ***P < 0.001; ****P < 0.0001; Mann–Whitney test).

Persistence of DNA damage can be passed to the next generation and be visible as “scars” in terms of high 53BP1 nuclear bodies (53BP1 NBs). Thus, we evaluated the presence of 53BP1 NBs in U2OS cells recovering from perturbed replication after RAD52 inhibition or combined POLA1i and RAD52i treatment. As shown in Figure 8B, replication perturbation in the presence of the RAD52i led to some increase in the number of 53BP1 NBs as compared with non-inhibited cells. However, combined treatment with RAD52i and POLA1i, significantly increased the fraction of cells with the higher number of 53BP1 NBs. Consistent with the increase in γH2AX-positive cells, the interference with the RAD51/Polα-mediated repriming induced in RAD52-inhibited cells by the low dose of POLA1i during treatment with 0.5mM HU resulted in a mild increase in total chromosomal damage but a larger stimulation of chromosomal fusions and exchanges (Supplementary Figure 14A-D).

Given that abrogation of the RAD51/Polα-mediated repriming pathway is critical for reducing DNA damage and compensating for RAD52 loss, we investigated whether this pathway also supported cell viability under conditions of replication stress. To this end, cells were exposed to 0.5 mM HU either with or without the RAD52 inhibitor and at both low and fully inhibitory doses of the Polα inhibitor (Figure 8C). Viability, assessed by crystal violet assay, demonstrated that combining RAD52 and Polα inhibitors - regardless of HU treatment - significantly reduced cell viability, whereas the low dose of the Polα inhibitor alone did not affect wild-type cells (Figure 8C). Additionally, in cells treated concurrently with 0.5 mM HU, the RAD52 inhibitor, and the low-dose Polα inhibitor, viability was similar to that observed in wild-type cells exposed to the high-dose Polα inhibitor (Figure 8C).

To substantiate further that the RAD51-Polα pathway is required for the recovery from replication stress in the absence of RAD52, we analyzed viability under perturbed replication of RAD52-inhibited cells expressing the ΔNTD-POLA1 alone or in combination with the low dose of Polαi. For better evaluation of the effects of the combined impairment of RAD51-Polα and RAD52-dependent pathways, cells were treated with 0.125µM HU (Figure 8D). As shown in Figure 8D, neither the inhibition of RAD52 nor the expression of the ΔNTD-POLA1 mutant was sufficient alone to reduce viability under perturbed replication. However, combined inhibition of RAD52 and expression of the ΔNTD-POLA1 mutant to disrupt the RAD51-Polα pathway of repriming significantly reduced viability and the observed reduction was not further increased by treatment with the low dose of Polαi (Figure 8D). As expected, treatment with the higher dose of Polαi reduced viability independently of the RAD52 inhibition.

Collectively, these results indicate that the RAD51/Polα pathway is a salvage mechanism that protects cells, experiencing perturbed replication, in the presence of a non-functional RAD52 from the accumulation of chromosomal damage, the passage to daughter cells of a damaged genome and cell death.

## DISCUSSION

Here, we have identified a novel, Polα-dependent repriming pathway that is activated in human cells under replication stress upon the loss of RAD52 function. Activation of this pathway leads to the accumulation of parental ssDNA gaps and represents a cellular response to perturbed replication distinct from the well-known PrimPol-dependent repriming mechanism ^17,18,20,23,33,48^.

Multiple pathways, including replication fork reversal and homologous recombination, have evolved to ensure recovery after replication fork perturbation or DNA damage. In addition, human cells can perform repriming downstream a DNA lesion or secondary DNA structures occurring in the leading strand by using a specialised primase/polymerase called PrimPol ^22^. This is opposed to what happens in yeast where repriming on both strands is performed by Polα ^21,49^. In human cells, PrimPol recruitment and repriming are also stimulated in BRCA2-depleted cells as a consequence of defective handling of RFs and loss of correct RAD51 function ^17,50^. Our findings indicate that specific defects in RF metabolism result in the accumulation of daughter-strand gaps, independent of PrimPol activity. Indeed, although some PrimPol recruitment is observed in the absence of RAD52, the majority of daughter-strand gaps is mediated by Polα in an origin-independent manner excluding the involvement of de-novo origin firing. Although our experimental approaches cannot rule out that ssDNA and DNA gaps accumulate away from the perturbed forks and do not represent DNA gaps caused by repriming at replication forks, these approaches have been widely used to identify replication-related DNA gaps.

Our data indicate that this Polα-dependent repriming is a downstream consequence of specific fork processing events in RAD52-deficient cells. The recruitment of Polα is dependent on the fork remodelling enzyme SMARCAL1 ^9^. Inhibition or mutations of RAD52 impairing its ability to bind to replication forks stimulate SMARCAL1 recruitment in vitro and in the cell ^16^. This suggest that Polα recruitment and repriming occur on stalled forks that have been remodelled by SMARCAL1 and subsequently degraded as shown when RAD52 is inhibited ^15^.

This observation raises a critical question: why does extensive fork degradation trigger Polα-dependent repriming in RAD52-deficient cells but not in BRCA2-deficient cells where degradation also occurs? In BRCA2-deficient cells, the RF is eventually cleaved by the MUS81 endonuclease, channelling the fork into break-induced replication (BIR) for restart^39^. RAD52, however, is required for MUS81 endonuclease activity at the fork ^38^.

Consequently, in RAD52-deficient cells, forks degrade extensively, possibly even more extensively than in BRCA2-deficient cells, but are not cleaved ^15^. Supporting this hypothesis, when we prevent fork cleavage in BRCA2-deficient cells by depleting MUS81, we restore recruitment of Polα at parental ssDNA (Figure 3D). This suggests that it is the extended degradation of a non-cleaved RF that creates the specific condition for this novel Polα-dependent pathway.

The recruitment of Polα to these degraded forks is critically dependent on RAD51. We observed an increased interaction between Polα and RAD51 in RAD52-inhibited cells, and even a minor depletion of RAD51 was sufficient to abrogate Polα recruitment and ssDNA accumulation. This high sensitivity to RAD51 levels is characteristic of processes requiring extensive RAD51 nucleofilament formation, such as fork protection or strand invasion, rather than more localized functions ^14^. RAD51 is important for fork restart and has been previously shown to interact with Polα for its localisation at the fork ^41^. Furthermore, it has been reported that RAD51 and MRE11 can stimulate an origin-independent re-loading of the replisome at collapsed replication forks ^51^. Our findings implicate a recombination-dependent replication mechanism (RDR). This mechanism has been well characterized in yeast but only few data were available in human cells ^52–55^. We propose that after extensive degradation of the RF, the resulting 3’-ended ssDNA flap is used by RAD51 to invade the template strand. RAD51 nucleoprotein filaments would stimulate recruitment of Polα at the D-loop structure for priming and new DNA synthesis. This differentiates this recruitment from that described previously, which has been shown to happen before fork reversal to limit its excessive engagement or to assist replication of perturbed lagging strand ^41,56^. Consistent with our model, expression of a RAD51 mutant destabilising nucleoprotein filaments and impairing D-loop formation abrogates Polα recruitment and ssDNA accumulation in RAD52-inhibited cells. Moreover, our *in vitro* biochemical assays show that a RAD51 nucleofilament formed on ssDNA but not RPA can robustly stimulate the priming and primer extension activity of the Polα-primase complex. Significantly, RAD51 also promotes primer extension by Polα on a substrate that models a D-loop, providing direct evidence for its ability to facilitate Polα-dependent synthesis within a recombination intermediate.

The role for Polα in human RDR is novel, as DNA synthesis during RDR in yeast is primarily driven by Polδ ^57^. In yeast, mutations leading to extensive processing of RFs can stimulate either Polα repriming or template-switching and repriming post strand-invasion, which might be dependent on Polα/primase ^21,58^. Thus, our observations could support a similar mechanism in human cells.

Noteworthy, Polα accumulation and ssDNA formation in RAD52-inhibited cells are abrogated in the presence of a POLA1 mutant that lacks the N-terminal region implicated in the interaction with RAD51 ^41^. These findings strongly suggest that the RAD51/Polα interaction is functional to the observed repriming pathway and point against the simple effect of template “melting” induced by RAD51 nucleofilaments. Interestingly, it has been recently reported that also Polθ can be recruited at the lagging strand to fill-in template gaps in the absence of RAD51 ^56,59^. Thus, multiple mechanism might contribute to prevent excessive fork reversal and accumulation of template gaps in the lagging strand. Although RAD52 loss or inhibition leads to a striking fork degradation phenotype, it does not impair fork restart ^15^. While BRCA2-deficient cells are dependent on BIR and PrimPol for restart ^32,39^, our data reveal that RAD52-deficient cells rely on this newly described RAD51-Polα-dependent pathway. Interference with this pathway using low amounts of a Polα inhibitor or expression of ΔNTD-POLA1 mutant unable to interact with RAD51 reduces fork progression and viability of RAD52-inhibited cells, also protecting from DNA damage accumulation. This RDR-like mechanism therefore is another crucial backup pathway for fork restart when canonical pathways are unavailable. This backup pathway might also contribute to explain why RAD52 knockout in humans does not generate a severe phenotype ^11^.

Altogether, our findings can be summarized in the model proposed in Figure 9. Under normal conditions, RAD52 is recruited to replication forks where it acts as a gatekeeper to prevent excessive SMARCAL1-driven fork reversal ^15,16^. In the absence of RAD52, stalled forks undergo uncontrolled reversal and extensive MRE11-dependent degradation. Because RAD52 is also needed for MUS81-mediated cleavage, these degraded forks cannot be channeled into BIR. This non-cleaved, degraded fork structure becomes a substrate for RAD51. Alternatively, RAD51-dependent Polα recruitment might occur after the extensive degradation of the RF, beyond the branching point (Figure 9). RAD51 nucleoprotein filaments would be assembled at the exposed parental ssDNA bringing Polα to the template strand for repriming. In both cases, DNA gaps would be filled-in or repaired post-replicatively. Thus, RAD51-dependent recruitment of Polα allows repriming of DNA synthesis facilitating fork progression and limiting chromosome breakage and cell death under perturbed replication. Notably, while preventing PrimPol-dependent repriming in BRCA2-deficient cells reduces formation of DNA damage ^60^ the suppression of RAD51/Polα-dependent repriming counteracts DNA damage further differentiating the two pathways.

**Figure 9.**
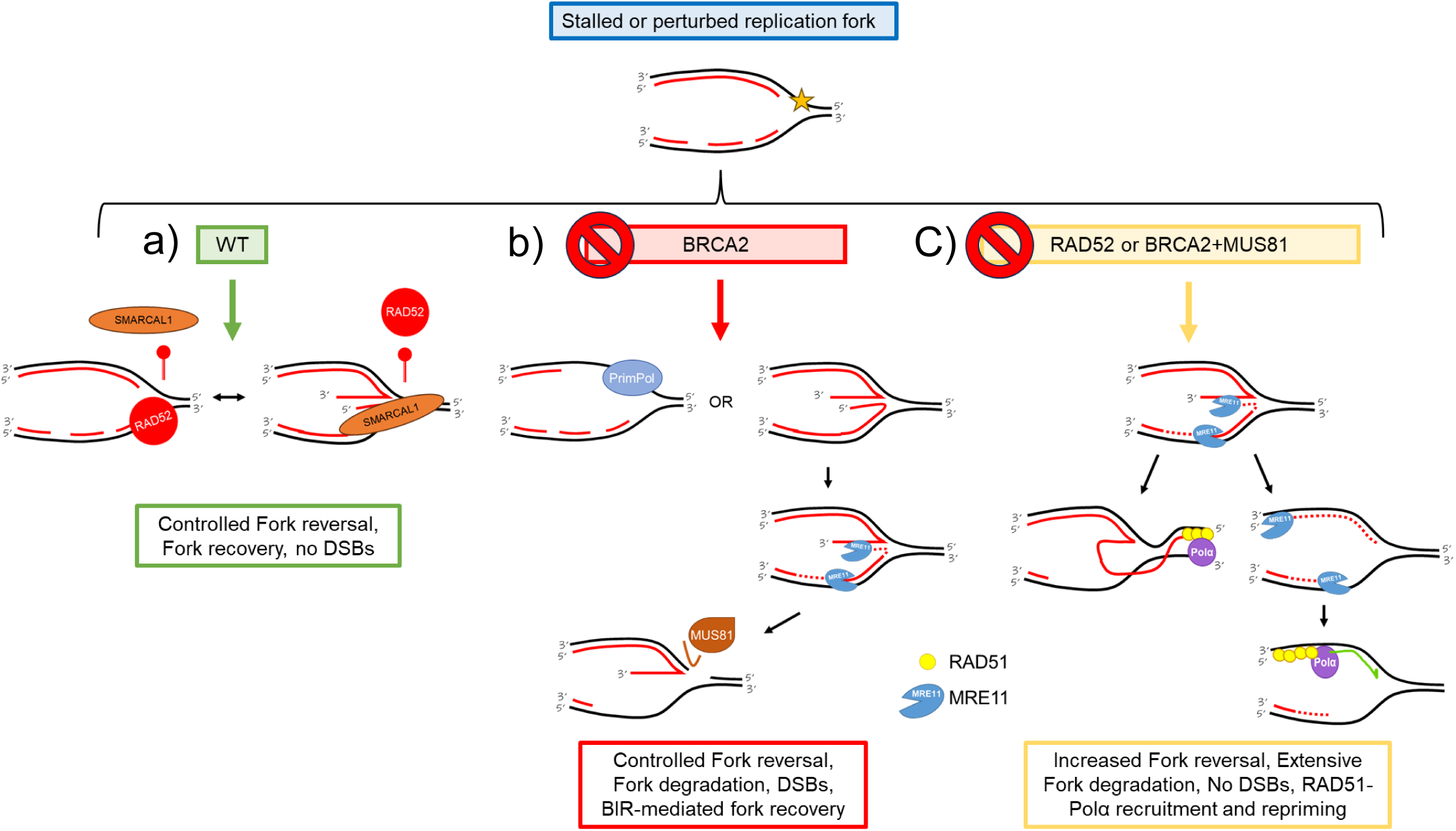
Model describing the RAD51-dependent Polα recruitment and gap formation downstream extensive degradation triggered by RAD52 deficiency. In response to perturbed replication, cells slow-down fork progression and engage fork remodelling factors such as SMARCAL1. RAD52 binds the forks to limit pathological fork reversal that could result in inefficient stabilisation of the forks once reversed (a). In the absence of BRCA2, the classic driver of fork de-protection, either repriming by PrimPol or degradation and breakage of the reversed forks takes place to promote restart (b). When RAD52 is absent or inhibited, or when MUS81 loss is added to BRCA2-depletion, perturbed replication forks undergo extensive reversal and prolonged exonucleolytic degradation leading to assembly of RAD51 nucleoprotein filaments at the parental ssDNA. Polα/primase is recruited by the interaction with RAD51 through its NTD region and stimulates repriming thus ensuring progression of perturbed forks (c).

As RAD52 is found mutated in cancer ^61^, it will be interesting to determine if these cancer-related mutations affect the gatekeeper role of RAD52 potentially rendering these tumours vulnerable to inhibitors of Polα. Moreover, RAD52 loss or inhibition is extremely synthetic sick with loss of BRCA2 or the checkpoint ^38,62,63^, and separation-of-function variants would be a useful tool to further characterize if this genetic relationship also correlates with an inability to perform RAD51/Polα-mediated repriming at distressed forks.

## MATERIALS AND METHODS

### Cell lines and culture conditions

The MRC5SV40 and human osteosarcoma cell line (U2OS) were maintained in Dulbecco’s modified Eagle’s medium (DMEM, Euroclone) supplemented with 10% fetal bovine serum (Euroclone) and incubated at 37 °C in a humidified 5% CO2 atmosphere. MRC5SV40 shRAD52 were obtained by transfection with a lentivirus expressing two different shRNA sequences (Sigma-Aldrich Mission lentivirus, sequence codes 271352 (V2)). MRC5SV40 KO PrimPol cells were a kind gift from Professor Aidan Doherty of Sussex University. MRC5SV40 KO MUS81 cells were obtained with the Crispr/Cas9 system by using two Alt-R® Crispr-Cas9 gRNA (Hs.Cas9.MUS81.1.AA and Hs.Cas9.MUS81.1.AB) and successively transfected with the WT protein ^40^. MRC5SV40 shBRCA2 or U2OS shSMARCAL1 were transfected with lentiviruses expressing the shSMARCAL1 or shBRCA2 cassette under the control of a doxycycline (Dox)-regulated promoter, at 0.5 of multiplicity of infection (MOI) (Dharmacon SmartVector inducible lentivirus, sequence code: V3SH11252-227970177 (shSMARCAL) and V3SH7669-226099147 (shBRCA2). After puromycin selection at 300 ng/ml, a single clone was selected and used throughout the study. MRC5SV40 ΔNTD-POLA1 cells were obtained by integrating the ΔNTD-POLA1 expression cassette into the AAVS1 site using the Crispr/Cas9 system and transfecting the two Vectorbuilder plasmids pRP[CRISPR]-hCas9-U6>AAVS1[gRNA#1] and VB231218-1466jab in a 1:1 ratio using the Neon transfection system (Invitrogen). After selection with neomycin and blasticidin, clones were obtained and one of retained for subsequent functional analyses.

Cell lines were routinely tested for mycoplasma contamination and maintained in cultures for no more than one month.

### Oligos and plasmids

#### siRNA

|  |  |
| --- | --- |
| MUS81 | Qiagen FlexiTube siRNA cat # SI04300877 oligo #6 |
| BRCA2 | Dharmacon ON TARGET PLUS SMARTpool siRNA #SO-2994934G |
| POLA1 | Designed by IDT: hs.Ri.POLA1.13.3 -SEQ1/2 |

#### Plasmids

|  |  |
| --- | --- |
| peGFP-PrimPol WT | Bailey et al., 2019 |
| pFLAG-MUS81 WT | Palma et al., 2018 |
| pRAD51-T131P | Liu et al., 2023 ([science.org/doi/10.1126/science.add7328](https://doi.org/10.1126/science.add7328)) |

### Transfections

All the siRNAs were transfected using Interferin® (Polyplus) 48 h before to perform experiments. MUS81 and BRCA2 siRNAs were used at 25 nM. POLA1-3’UTR siRNA was used at 40nM. The peGFP-PrimPol and plasmid expressing RAD51 isoforms were transfected in cell lines using Neon transfection system (Invitrogen) 48 h prior to perform experiments.

### Chemicals

HU (Sigma-Aldrich) was added to culture medium at 2 mM or 0.5 mM from a 200 mM stock solution prepared in Phosphate-buffer saline solution (PBS 1X) to induce DNA replication arrest or slowing. RAD52 inhibitor, Epigallocatechin (EGC - Sigma-Aldrich) was dissolved in DMSO at 100 mM and used at 50 µM. The B02 compound (Selleck), an inhibitor of RAD51 activity, was used at 27 µM. CDC7i (XL413, Selleck) was dissolved in sterile water and used 10 µM. Doxycycline (Sigma-Aldrich) was dissolved in DMSO and used 1µ/ml. MIRIN, the inhibitor of MRE11 exonuclease activity (Calbiochem), was used at 50 µM. Polα inhibitor (ST1926 – Sigma-Aldrich) was dissolved in DMSO and used at the concentrations reported on the experiments. S1 nuclease (Invitrogen cat # 18001016) was diluted 1/100 in S1 buffer and used at 20 U/ml. CldU (Sigma-Aldrich) was dissolved in sterile water as a 200 mM stock solution and used at 50 μM. IdU (Sigma-Aldrich) was dissolved in sterile DMEM as a stock solution 2.5 mM and used at 100 for ssDNA and PLA assays or 250 μM for fibers assay. 5-ethynyl-2’-deoxyuridine (EdU) (Sigma-Aldrich) Was dissolved in DMSO and used at 10 μM for EdU incorporation assay or 100 μM for SIRF assay.

### Western blot analysis

Western blots were performed using standard methods. Blots were incubated with primary antibodies against: anti-GFP (Santa Cruz Biotechnology, 1:500), anti-MUS81 (Santa Cruz Biotechnology, 1:1000), anti-LAMIN B1 (Abcam, 1:10,000), anti-GAPDH (Millipore, 1:5000), anti-RAD51 (Abcam 1:10,000), anti-SMARCAL1 (Bethyl 1:1500), anti-PrimPol (1:1000 Proteintech), anti-BRCA2 (Bethyl 1:5000), anti-POLA1 (Polα - Bethyl, 1:500). After incubations with horseradish peroxidase-linked secondary antibodies (Jackson ImmunoResearch, 1:20,000), the blots were developed using the chemiluminescence detection kit ECL-Plus (Amersham) according to the manufacturer’s instructions. Quantification was performed on blot acquired by ChemiDoc XRS+ (Bio-Rad) using Image Lab software, the normalization of the protein content was done through LAMIN B1 or GAPDH immunoblotting.

### Chromatin isolation

After the treatments, cells (4 × 106 cells/ml) were resuspended in buffer A (10 mM HEPES, [pH 7.9], 10 mM KCl, 1.5 mM MgCl2, 0.34 M sucrose, 10% glycerol, 1 mM DTT, 50 mM sodium fluoride, protease inhibitors [Roche]). Then Triton X-100 was added at a final concentration of 0.1% and the cells were incubated for 5 min on ice. Nuclei were collected in pellets by low-speed centrifugation (4 min, 1300 × g, 4 °C) and washed once in buffer A. Nuclei were then lysed in buffer B (3 mM EDTA, 0.2 mM EGTA, 1 mM DTT, protease inhibitors) for 10 min on ice. Insoluble chromatin was collected by centrifugation (4 min, 1700 × g, 4 °C), washed once in buffer B + 50 mM NaCl, and centrifuged again under the same conditions. The final chromatin pellet was resuspended in 2X Laemmli buffer and sonicated for 15 s in a Tekmar CV26 sonicator using a microtip at 50% amplitude.

### EdU incorporation assay

U2OS were treated with 10 µM EdU 10 min before giving the treatment with ST1926 for 30 min at the indicated concentrations. After the treatment, cells were permeabilized with 0.5% TritonX-100 in PBS 1X for 10 minutes on ice, then fixed with 3% PFA, 2% sucrose in PBS 1X at room temperature (RT) for 15 min. For the EdU detection was applied the Click-iT™ EdU Alexa Fluor™ 488 Imaging Kit (Invitrogen) for 30 minutes at RT. The reagents for the Click-iT™ reaction were diluted according to the manufacturer’s instructions. Nuclei were examined with Eclipse 80i Nikon Fluorescence Microscope, equipped with a Virtual Confocal (ViCo) system and foci were scored at 40X magnification. Quantification was carried out using the ImageJ software.

### IdU incorporation assay

MRC5SV40 and U2OS were treated with 100 µM IdU for 20 hours and released for 2 hours in fresh DMEM before giving the treatment with RAD52i and HU for 4 hours at the reported concentrations. After the treatment, cells were fixed at RT with 4% PFA/PBS for 10 min, permeabilized with 0.4% TritonX-100/PBS and denatured with 2.5 N HCl/PBS for 45 minutes. Cells were then incubated with mouse anti-IdU antibody (Becton Dickinson, 1:80) for 1 h at 37 °C in 1% BSA/PBS, followed by species-specific fluorescein-conjugated secondary antibodies (Alexa Fluor 488 Goat Anti-Mouse IgG (H + L), highly cross-adsorbed—Life Technologies). Slides were analysed with Eclipse 80i Nikon Fluorescence Microscope, equipped with a Virtual Confocal (ViCo) system. Quantification was carried out using the ImageJ software.

### Detection of ssDNA by native IdU assay

To detect parental ssDNA, cells were labelled for 20 hours with 50 µM IdU (Sigma-Aldrich), released in fresh DMEM for 2 hours, then treated as indicated. For immunofluorescence, cells were washed with PBS 1X, permeabilized with 0.5% Triton X-100 for 10 min at 4 °C and fixed in 3% PFA, 2% sucrose in PBS 1X. Fixed cells were then incubated with mouse anti-IdU antibody (Becton Dickinson, 1:80) for 1 h at 37 °C in 1% BSA/PBS, followed by species-specific fluorescein-conjugated secondary antibodies (Alexa Fluor 488 Goat Anti-Mouse IgG (H + L), highly cross-adsorbed—Life Technologies). Slides were analysed with Eclipse 80i Nikon Fluorescence Microscope, equipped with a Virtual Confocal (ViCo) system. For each point, at least 100 nuclei were analysed. Quantification was carried out using the ImageJ software.

### BrdU alkaline Comet assay

For the BrdU alkaline Comet assay, cells were incubated with 100 μM BrdU and 0,5Mm of HU for 1 hour, then cells were washed with PBS 1X followed by incubation in fresh media for 1 hour. Cells were harvested in PBS, kept on ice to inhibit DNA repair and then subjected to the alkaline Comet assay. Dustfree frosted-end microscope slides were then dipped into molten agarose at 1% and left to dry. Cell suspensions were rapidly mixed with LMP agarose at 0.5% kept at 37°C and an aliquot was pipetted onto agarose-covered surface of the slide. Agarose embedded cells were lysed by submerging slides in lysis solution (NaCl 2,5M; EDTA 100mM; Trizma 10mM; 0.2N NaOH; 1% Triton X-100; 10% DMSO) and incubated at 4°C, overnight in the dark. After lysis, slides were washed in running buffer pH13 (NaOH 10N; EDTA 200 mM) for 1 min. Electrophoresis was performed for 20 min in running buffer pH13 at 25 V/250-300 mA. Slides were subsequently washed in distilled H2O and finally dehydrated in ice cold methanol. For the immunofluorescence, slides were stained with primary mouse anti-IdU antibody (Becton Dickinson, 1:80) for 1 h at 37 °C in 1% BSA /PBS, followed by species-specific fluorescein-conjugated secondary antibodies (Alexa Fluor 488 Goat Anti-Mouse IgG (H + L), highly cross-adsorbed—Life Technologies). Slides were analysed with Eclipse 80i Nikon Fluorescence Microscope, equipped with a Virtual Confocal (ViCo) system. For each time point, at least 50 nuclei from triplicate experiments were analysed. The olive tail moment was calculated using CometScore 2.0 software.

### In situ PLA assay for ssDNA–protein interaction

The in-situ PLA NaveniFlex (Navinci Diagnostics) was performed according to the manufacturer’s instructions. For parental ssDNA-protein interaction, cells were labelled with 100 μM IdU for 20 hours and then released in fresh medium for 2 hours. After treatment, cells were permeabilized with 0.5% Triton X-100 for 10 min at 4 °C, fixed with 3% PFA/2% sucrose in PBS 1X for 10 min and then blocked in 3% BSA/PBS for 15 min. After washing with PBS, cells were incubated with the two relevant primary antibodies. The primary antibodies used were as follows: mouse monoclonal anti-RAD51 (GeneTex, 1:150), mouse monoclonal anti-IdU (Becton Dickinson, 1:50), rabbit polyclonal anti-Polα (Bioss, 1:50), rabbit polyclonal anti-GFP (Invitrogen 1:150), rabbit anti-WDHD1 (Novus Biologicals, 1:50), rabbit anti-RAD52 (Aviva, 1:150). The negative controls were obtained by using only one primary antibody. Samples were incubated with secondary antibodies conjugated with PLA probes MINUS and PLUS: the PLA probe anti-mouse PLUS and anti-rabbit MINUS (NaveniFlex equivalent). The incubation with all antibodies was accomplished in a humidified chamber for 1 h at 37 °C. Next, the PLA probes MINUS and PLUS were ligated using two connecting oligonucleotides to produce a template for rolling-cycle amplification. After amplification, the products were hybridized with red fluorescence-labelled oligonucleotide. Samples were mounted in Prolong Gold anti-fade reagent with DAPI (blue). Images were acquired randomly using Eclipse 80i Nikon Fluorescence Microscope, equipped with a Virtual Confocal (ViCo) system. The analysis was carried out by counting the PLA spot for each nucleus. Analysis of PLA spots was performed by two independent investigators by zooming out each nucleus and performing manual counting. Nuclei showing uncountable spots even in magnified images where not analysed. To exclude the cells outside the S-phase from the analysis, nuclei with less than 7 (treated) or 2 spots (untreated) were considered as negative.

### Single-cell Assay for in situ Protein Interaction with Nascent DNA (SIRF)

Exponential growing cells were seeded onto microscope chamber slide. On the day of experiment, cells were incubated with 100 µM EdU for 20 min and treated as indicated. After treatment, cells were pre-extracted in 0.5% TritonX-100 for 5 min on ice and fixed with 3% PFA, 2% sucrose in PBS 1X for 15 min at RT. Cells were then blocked in 3% BSA/PBS for 15 min. For the EdU detection was applied the Click-iT™ EdU Alexa Fluor™ Imaging Kit (Invitrogen) using 5mM Biotin-Azide for 30 minutes at RT. The primary antibodies used were as follows: mouse monoclonal anti-RAD51 (GeneTex, 1:150), rabbit polyclonal anti-Polα (Bioss, 1:50), mouse anti-biotin (Invitrogen, 1:50) rabbit anti-biotin (Abcam, 1:50). The negative controls were obtained by using only one primary antibody. Samples were incubated with secondary antibodies conjugated with PLA probes MINUS and PLUS: the PLA probe anti-mouse PLUS and anti-rabbit MINUS (Duolink®, Sigma-Aldrich). The incubation with all antibodies was accomplished in a humidified chamber for 1 h at 37 °C. Next, the PLA probes MINUS and PLUS were ligated using two connecting oligonucleotides to produce a template for rolling-cycle amplification. After amplification, the products were hybridized with red fluorescence-labelled oligonucleotide. Samples were mounted in Prolong Gold antifade reagent with DAPI (blue). Images were acquired randomly using Eclipse 80i Nikon Fluorescence Microscope, equipped with a Virtual Confocal (ViCo) system. The analysis was carried out by counting the SIRF spot for each nucleus.

### Single-Molecule Localisation Microscopy

U2OS cells were treated as indicated in the experimental scheme. After the treatment, cells were fixed and immunofluorescence was performed as described by Whelan & Rothenberg, 2021. EdU was detected with the Click-iT™ EdU Alexa Fluor™ Imaging Kit (Invitrogen) for 30 minutes at RT. Coverslips with fixed cells were stored for up to 1 week at 4 °C prior to dSTORM imaging. The fixed cells on coverslips were mounted onto concave slides and dSTORM imaging B3 buffer from Oxford Nanoimaging (ONI, PN #900-00004) was added before imaging. A commercial TIRF microscope (Nanoimager S Mark IIB from ONI, https://oni.bio) with lasers of 405 nm/150 mW, 488 nm/1 W, 561 nm/1 W, and 640 nm/1 W was used to acquire data. Briefly, fluorophore emission was collected by an oil immersion 100x/1.45NA objective and images were acquired at 30 ms exposure time using a Hamamatsu ORCA-Flash4.0 V3 Digital sCMOS camera. The reconstruction of the super-resolution image was conducted by NimOS software from ONI. Localization data were then analysed using the ONI-CODI platform for drift correction, filtering, clustering, and counting.

### Protein Preparation

Homogeneity human protein sample preparation of Polα/primase ^64^, RPA ^65^ and RAD51 ^66^ have been described. Site-specific F86E and T131P mutations were introduced in the RAD51 genes in the plasmids by Genscript mutagenesis service (Genscript, Piscataway, NJ) and corresponding RAD51 mutant protein were prepared as wild type.

### Primase Assay

The priming activity of human Polα/primase was tested on poly-dT70 (IDT, Coralville, IA). Indicated amount of Polα/primase and ssDNA templates (500 nM) were incubated at 30 °C for 60 min in a standard reaction (20 μl) consisted of 20 mM Tris-Acetate (pH 8.0), 50 mM NaCl, 50 mM NaCl, 4 mM MgCl_2_, 1 mM spermidine, 5 mM DTT, 0.5 μM [α-^32^P] ATP (3000 Ci/mmol^−1^, Revvity), 50 μM unlabelled ATP and 4 U of RiboLock RNase Inhibitor (Thermo Scientific). The indicated amount of RAD51 or RPA was added into the reaction and held for 5 min before adding Polα/primase. Reaction products were mixed with 30 μl formamide loading buffer (90% formamide, 30 mM EDTA, 0.5× TBE, 0.1% of bromophenol and 0.025% SDS) and heated at 80 °C for 5 min. Samples (10 μl) were loaded on a sequencing-style 15% acrylamide, 7 M urea, 1× TBE gel. The bromophenol blue dye was run to the bottom of the gel (90 min, 250V). Radiolabeled RNA synthesis products were imaged on a FLA-7000 image scanner (FUJIFILM). Unless indicated otherwise, Polα–primase activity was determined by total counts per spot or relative product (%) in which 100% indicate total counts of Polα/Primase only products.

### Primer Extension Assay

RNA primer extension activities of the Polα/primase in various conditions were compared in reactions (20 μl) that contained the 0.5 μM poly-dT_70_ template with Cy3 fluorophore-labeled poly-rA_15_ RNA oligos. The annealing of the template with RNA primers was done by a decrease of temperature from 90 to 25 °C in 0.2 °C/min gradient in the thermo cycler. Reactions were assembled on ice in the following order. The primer/template was added to the buffer containing 20 mM Tris-HCl (pH 7.5), 50 mM NaCl, 10 mM MgCl_2_, 0.2 mg/ml BSA, and 2 mM DTT followed by the addition of dATP (0.2 mM). Then, the indicated amount of RAD51 or RPA was added into the reaction and held for 5 min before Polα/primase was added and further incubated for 30 min at 30 °C. Reaction products were separated as described in primase assay. Visualization of the products used the Chemidoc MP imager (Bio-Rad).

### Electrophoresis Mobility Shift Assay

Indicated concentrations of RAD51 or RPA were mixed with 500 nM of 5’-end Cy3 labeled poly-dT70 DNA substrates in a 15 µl of standard priming reaction buffer. The reaction mixtures were incubated at 30°C for 5 min then 1.5 µl of 10x Orange-G loading dye [0.1% Orange-G, 50% glycerol, 400 mM Tris-Acetate (pH 8.0) and 10 mM EDTA] was added. The bound and free DNA species were resolved by the non-denaturing 5.5% polyacrylamide gel electrophoresis in TAE buffer [40 mM Tris-Acetate (pH 8.0) and 1 mM EDTA]. The gel was imaged using a ChemiDoc MP (Bio-Rad) by exciting and monitoring Cy3 fluorescence.

### DNA fibers analysis

Fiber’s assay using S1 nuclease was performed as indicated by Quinet et al., 2017. Briefly, cells were pulse-labelled with 50 µM CldU and then labelled with 250 µM IdU with or without treatment, as reported in the experimental schemes. At end of treatment, cells were permeabilized with CSK buffer (100 mM NaCl, 10 mM PIPES pH 6.8, 1M EGTA, 3 mM MgCl2, 300 mM sucrose, 0.5% Triton X-100) for 10 min at RT, then were washed with PBS 1X and S1 nuclease buffer (30 mM sodium acetate, 10 mM zinc acetate, 5% glycerol, 50 mM NaCl) prior to add +/- S1 nuclease for 30 min at 37 °C in a humidified chamber. Cells were washed with S1 buffer then scraped with 0.1% BSA/PBS and collected pellets were used to perform fibers spreading. For the fork progression assay, cells were pulse-labelled with 50 µM CldU for 20 min and then labelled with 250 µM IdU for 40 min, with or without RAD52i. DNA fibers were spread out as described ^27^. For immunodetection of labelled tracks, the following primary antibodies were used: rat anti-CldU/BrdU (Abcam 1:50) and mouse anti-IdU/BrdU (Becton Dickinson 1:10). Images were acquired randomly from fields with untangled fibers using Eclipse 80i Nikon Fluorescence Microscope, equipped with a Virtual Confocal (ViCo) system. The length of labelled tracks was measured using the Image-Pro-Plus 6.0 software.

### Viability assay

For treatments with the Polαi, cells were seeded in 48-well plates. Once attached, they were subjected to the treatments (inhibitors and HU at the concentrations indicated above). The cells remained in culture until the positive control (untreated WT) reached confluence (about a week), and the treated medium was changed every 3 days. The cells were then stained with 0.5% crystal violet, 25% methanol in double-distilled water for 24 hours. The wells were rinsed with double-distilled water and allowed to dry. Once completely dry, 200 µL of methanol was added to each well and left for 20 minutes under gentle agitation so that the methanol decolorized the cells at the bottom of the well and the color transferred into the solution. The medium was then transferred to 96 multiwell plates and the absorbance values are recorded at 570nm using a plate reader. The values (N≥3 were analysed using the Mann-Whitney test.

For cell proliferation, cells were harvested at 24-, 48- and 72-hours post-transfection, stained with trypan blue and counted in triplicate on the TC20 Automated Cell Counter.

### Chromosomal aberrations

MRC5SV40 cells were treated with HU 2 mM, with or without RAD52 inhibitor and/or Polα inhibitor, left at 37 °C for 4 hours, then allowed to recover in fresh medium for additional 24 hours. Cell cultures were incubated with colcemid (0.2 µg/ml) at 37 °C for 3 hours until harvesting. Cells for metaphase preparations were collected and prepared as previously reported ^67^. For each condition, at least 50 chromosomes were examined, and chromosomal damage scored at 100× magnification.

### Statistical analysis

Experiments shown are representative of at least two independent biological replicates unless otherwise indicated in the figure legend. Significance was assessed using the built-in tools in Prism 9 (GraphPad Inc.) by Kruskal-Wallis’ test followed by post-hoc Dunn test for FDR for experiments with more than two samples, and by the two-tailed Student’s t-test to evaluate the means from normal distributions when analysing two samples. P < 0.05 was considered as significant. Statistical significance was always denoted as follow: ns = not significant; *P < 0.05; **P < 0.01; ***P < 0.01; ***P < 0.001. If not otherwise indicated in the figure legend, analysis was performed by Kruskal-Wallis’ test followed by post-hoc Dunn. Specific statistical analyses are reported in the relevant legend. No statistical methods or criteria were used to estimate sample size or to include/exclude samples.

## ACKNOWLEDGMENTS

We thank Prof. Aidan Doherty for PrimPol KO cells and GFP-PrimPol expressing plasmid. We thank Prof. David Cortez for the RAD51-T131P and RAD51 wild-type expression plasmids. We thank Prof. C.J. Lim for the purified recombinant Polα/Primase complex and assistance with assay. We want to thank Luca Pellegrini for helpful discussion and advise on RAD51-NTD POLA1 interaction. This work was supported by NIH-NCI grant (grant n. R01CA232425-A1) to M.S. and P.P., and by investigator grants from Associazione Italiana per la Ricerca sul Cancro (AIRC) to P.P. (IG n. 28972) and to A.F. (IG n. 30446).

## AUTHORS CONTRIBUTION

G.M. performed analysis of ssDNA and protein recruitment by QIBC, generated the new cell models and performed experiments with ΔNTD-POLA1 and RAD51 mutants. L.D.B performed initial experiments to analyse formation and mechanism of Polα-dependent parental DNA gaps using RAD52i. E.M. performed analysis of parental ssDNA formation in RAD52 knockdown cells and contributed to fork recovery assay and ΔNTD-POLA1 characterisation. M.H. performed the in vitro biochemical experiments. G.M, E.M. and M.H. took care of all the additional experiments to identify the relationship between RAD51 and Polα. P.V. performed the analyses of chromosome damage. E.M. and F.A.A. performed dSTORM experiments. All authors contributed to design experiments and analysed data. P.P, A.F and M.S. designed experiments, analysed data and supervised the project. All authors contributed to revise the paper.

