## Supplementary Figures and Legends for "RAD52 prevents accumulation of RAD51 and Polα-dependent DNA gaps at perturbed replication forks"

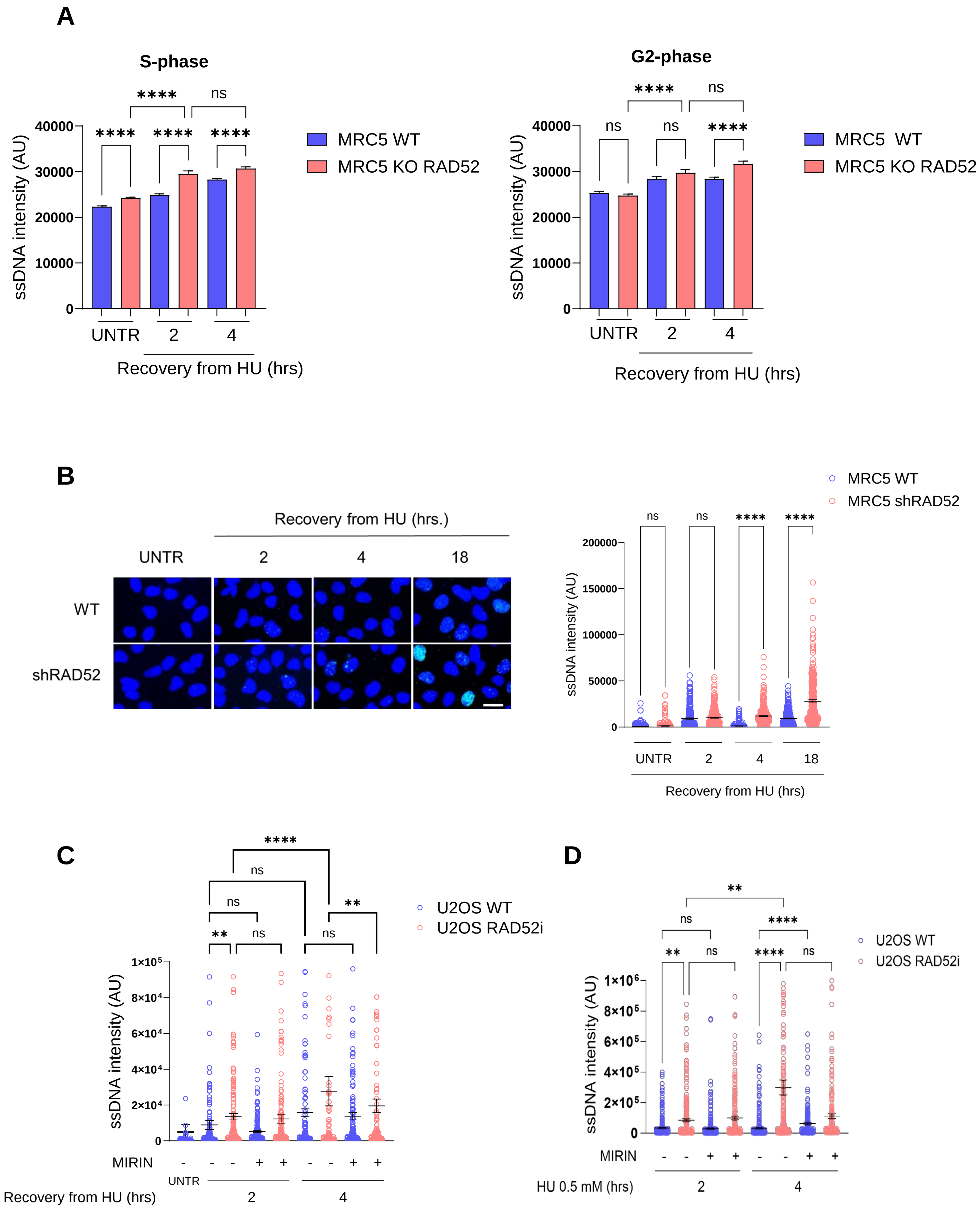


Supplementary Figure 1

A

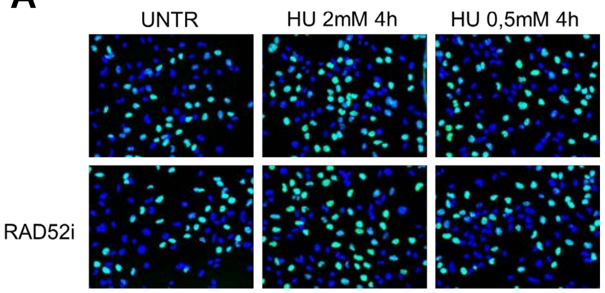


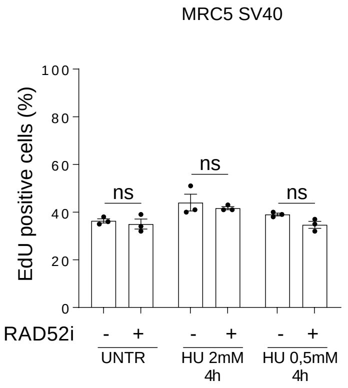


B

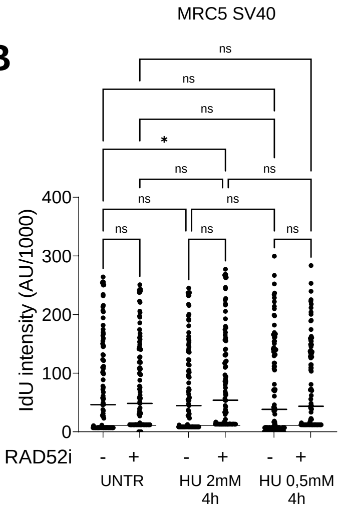


C

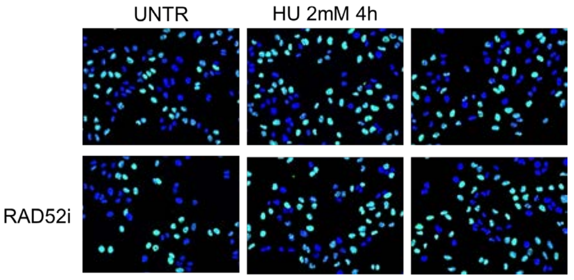


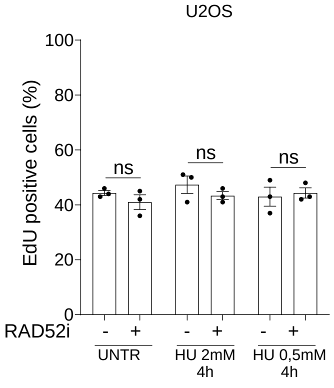


Supplementary Figure 2

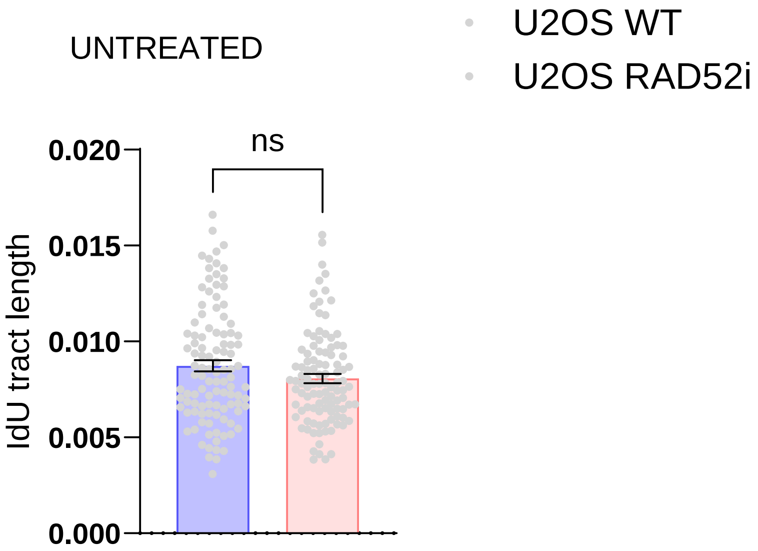


Supplementary Figure 3

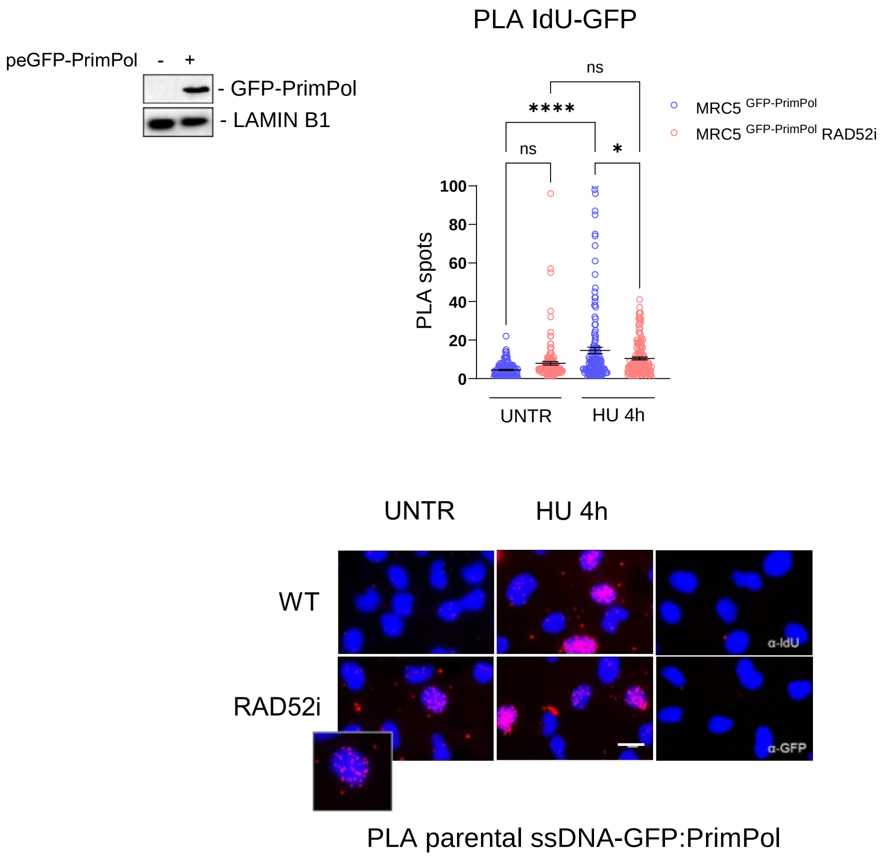


Supplementary Figure 4

**A**

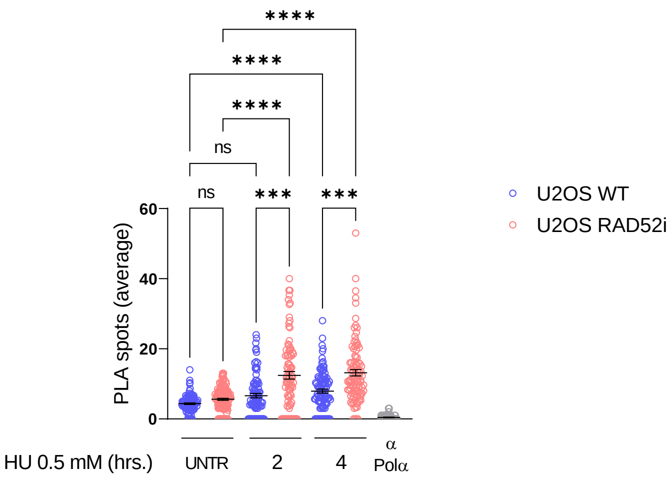


**B**

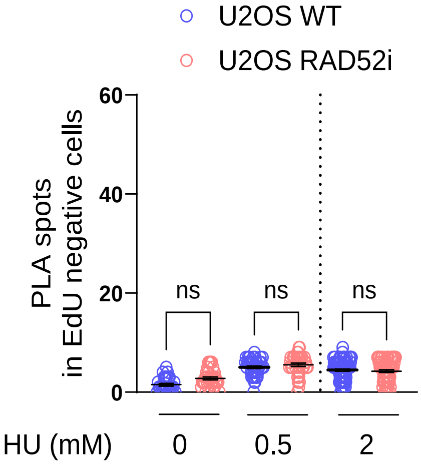


Supplementary Figure 5

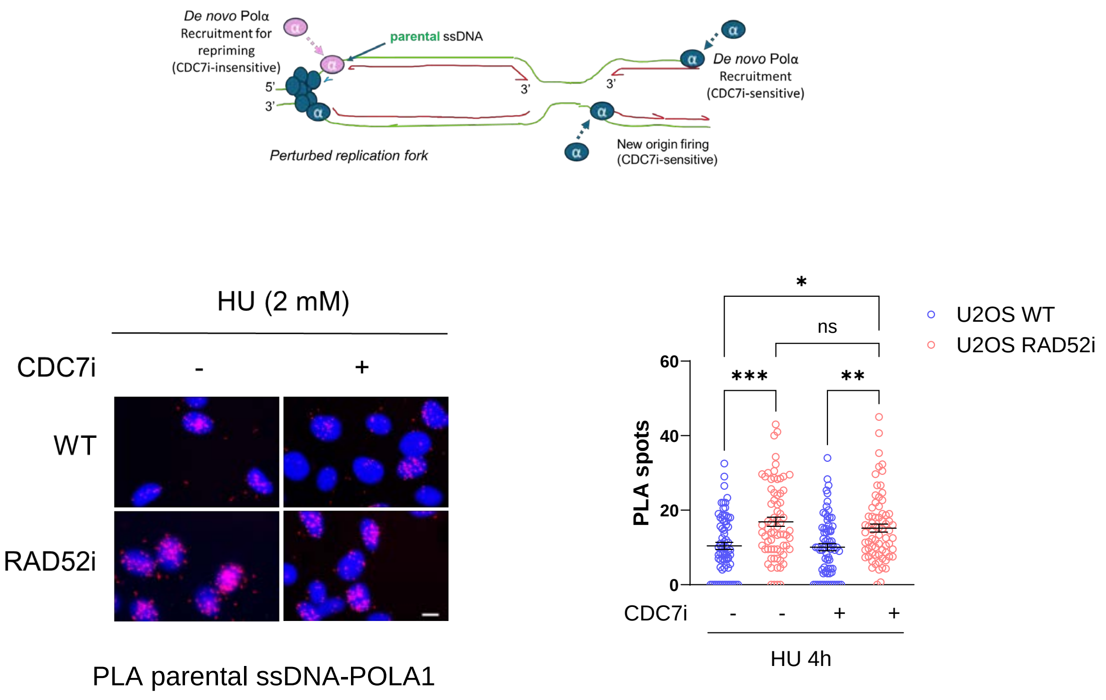


Supplementary Figure 6

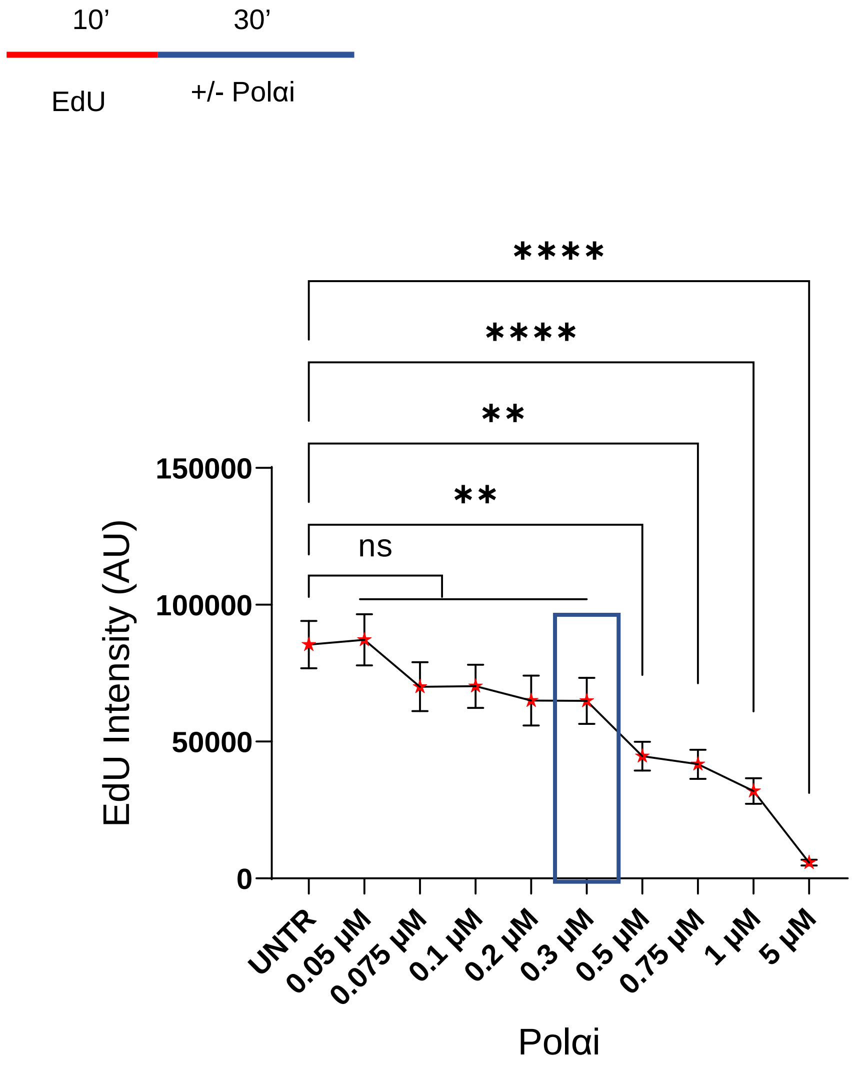


Supplementary Figure 7

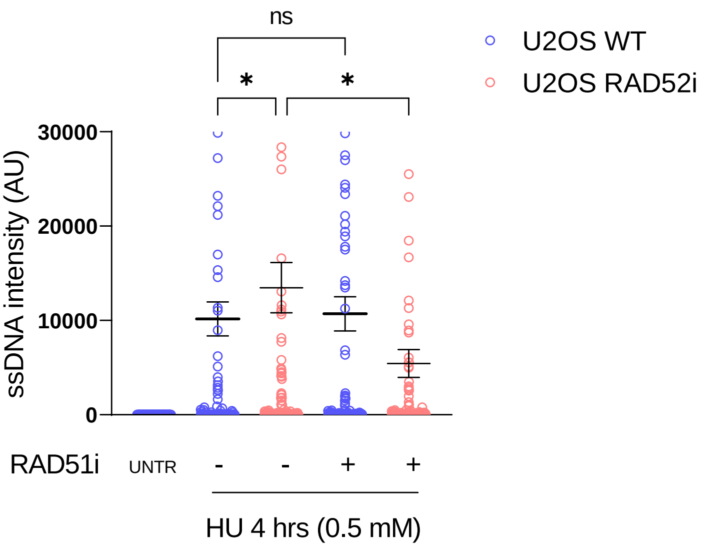


Supplementary Figure 8

**A**

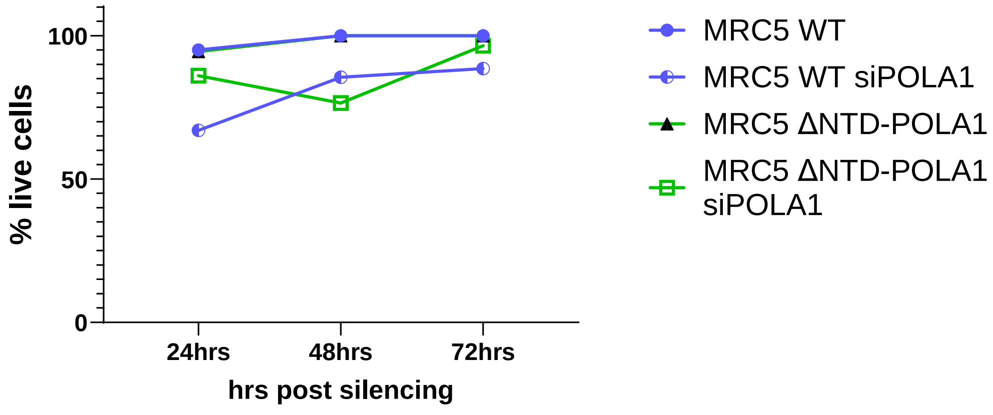


**B**

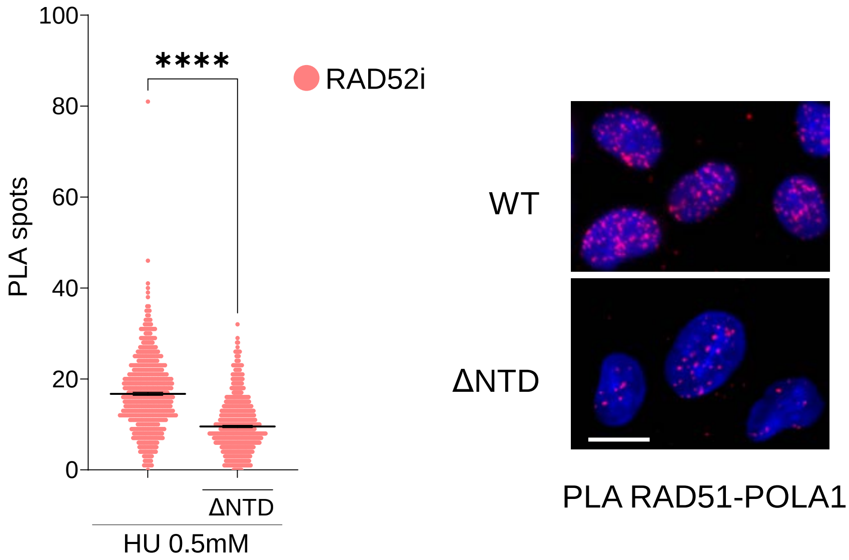


Supplementary Figure 9

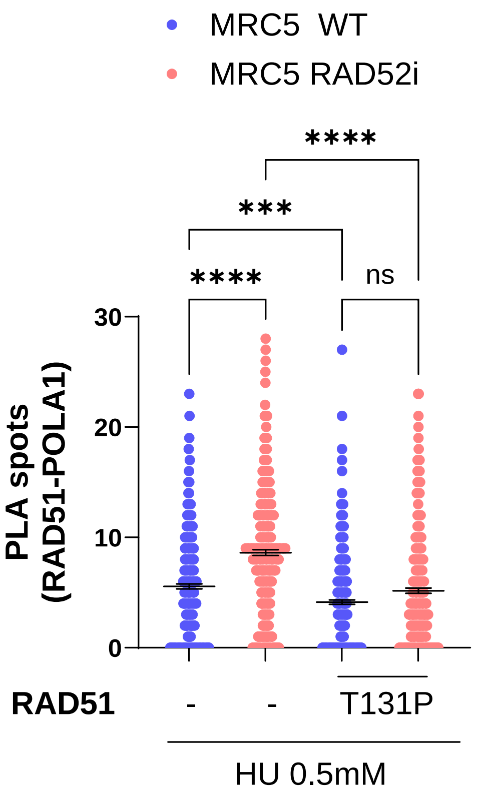


Supplementary Figure 10

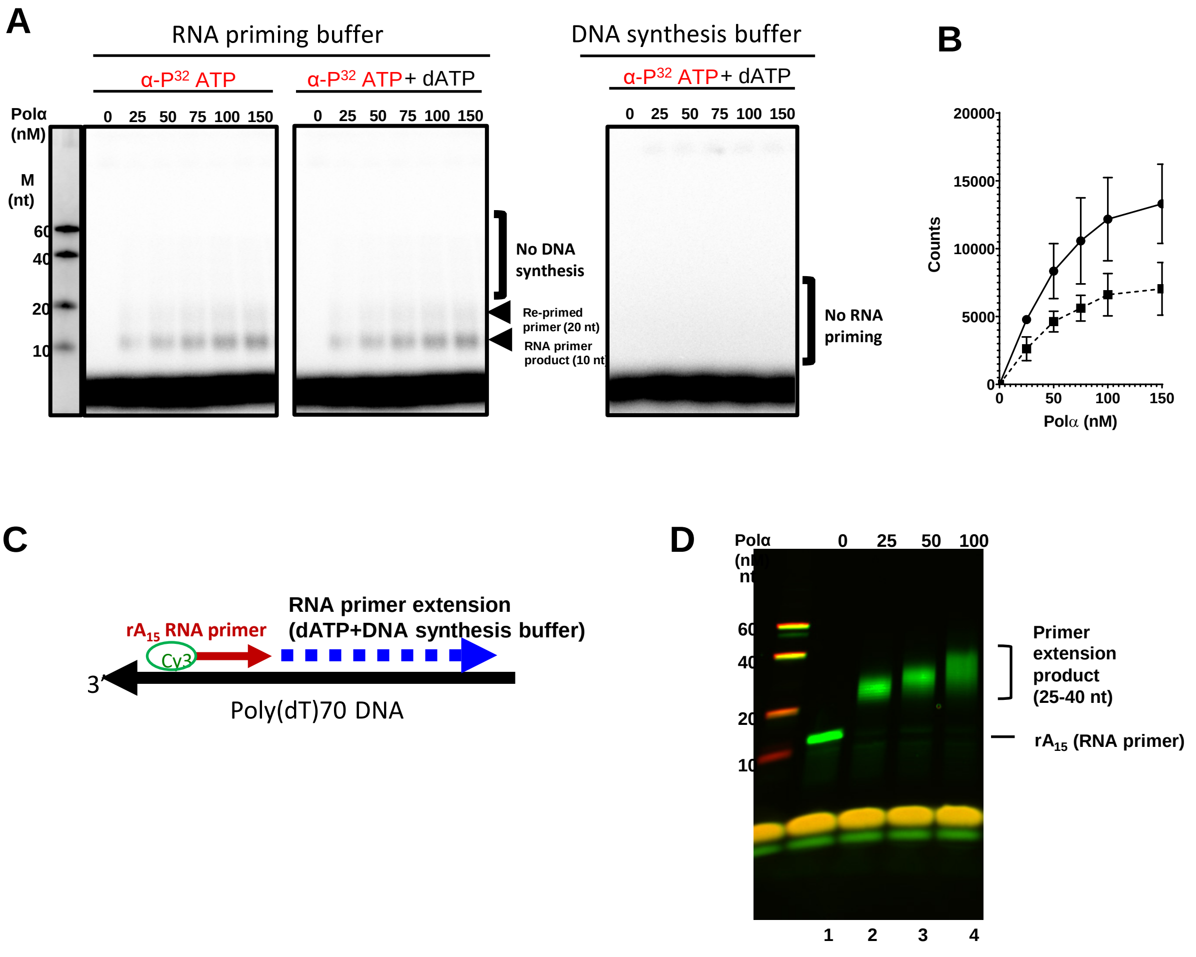


Supplementary Figure 11

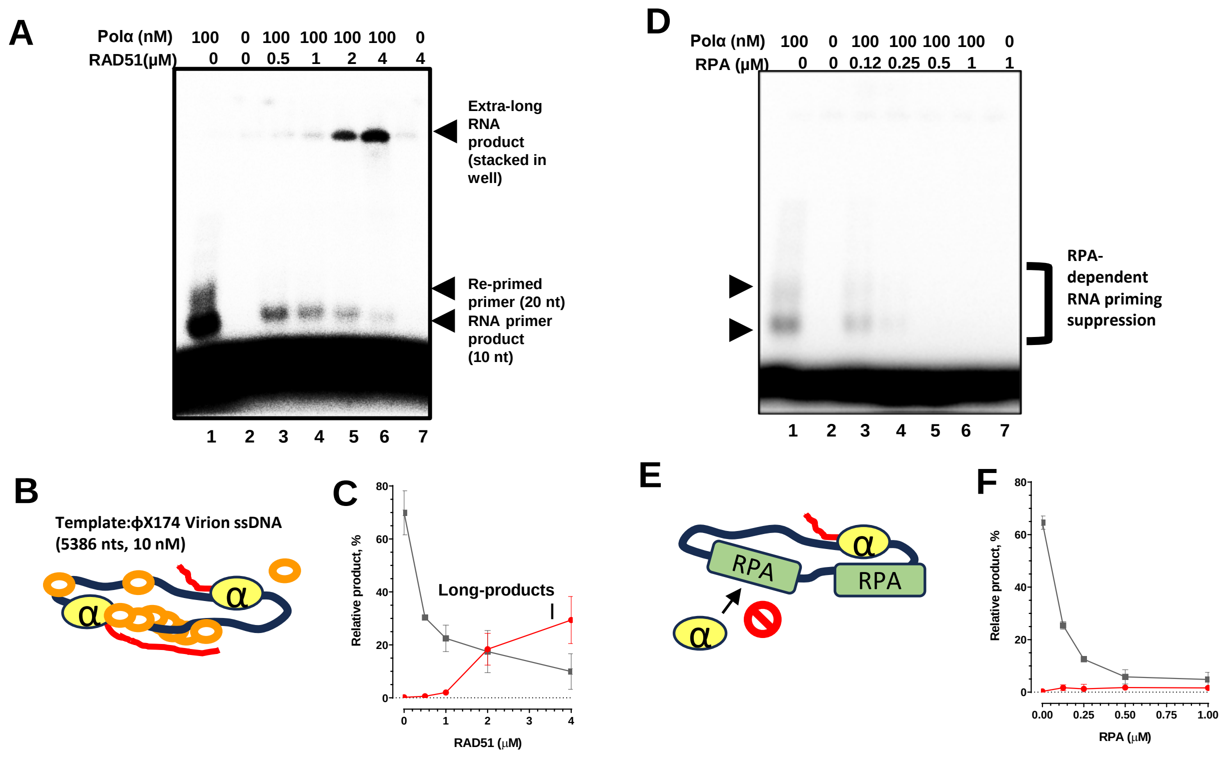


Supplementary Figure 12

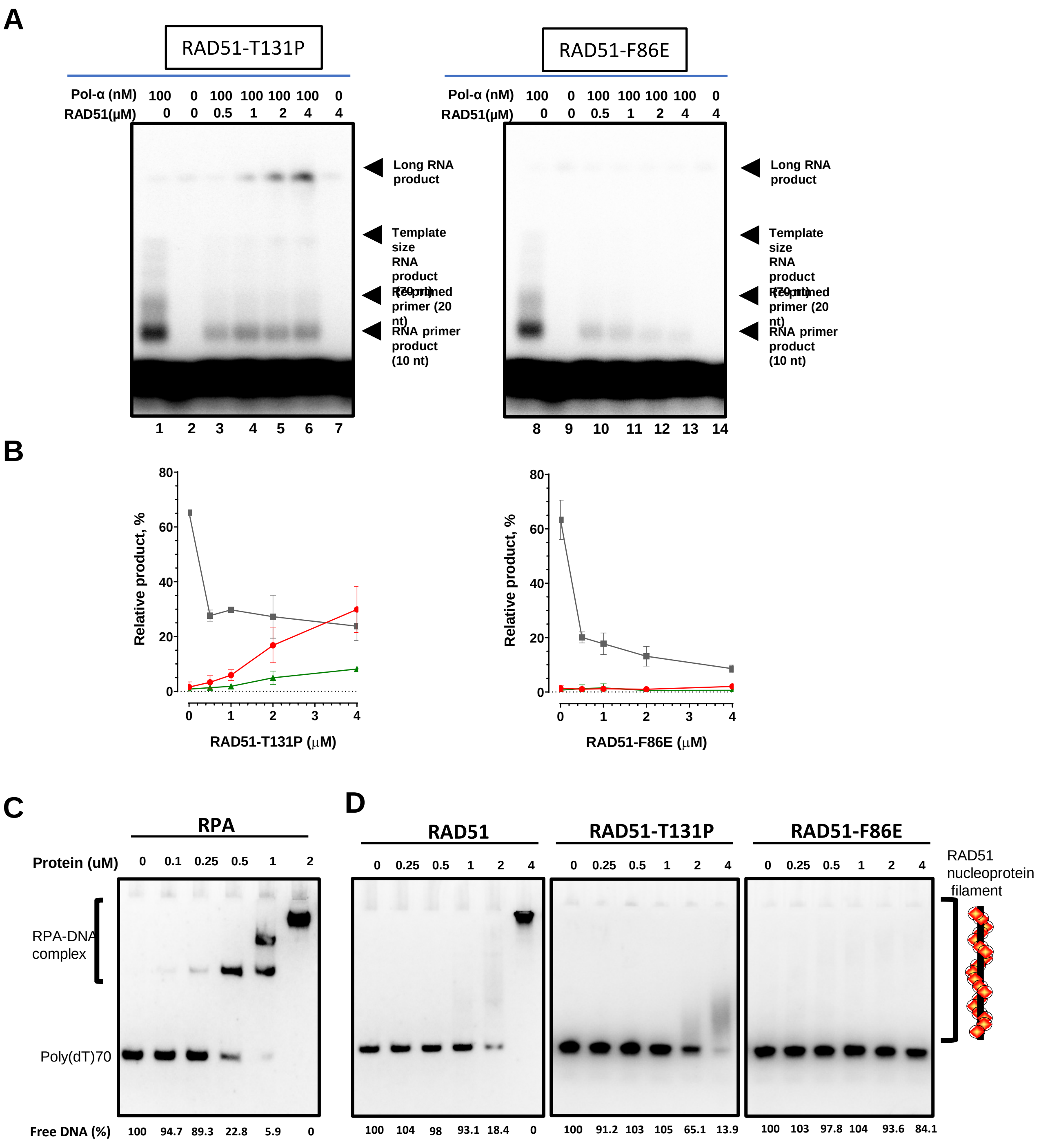


Supplementary Figure 13

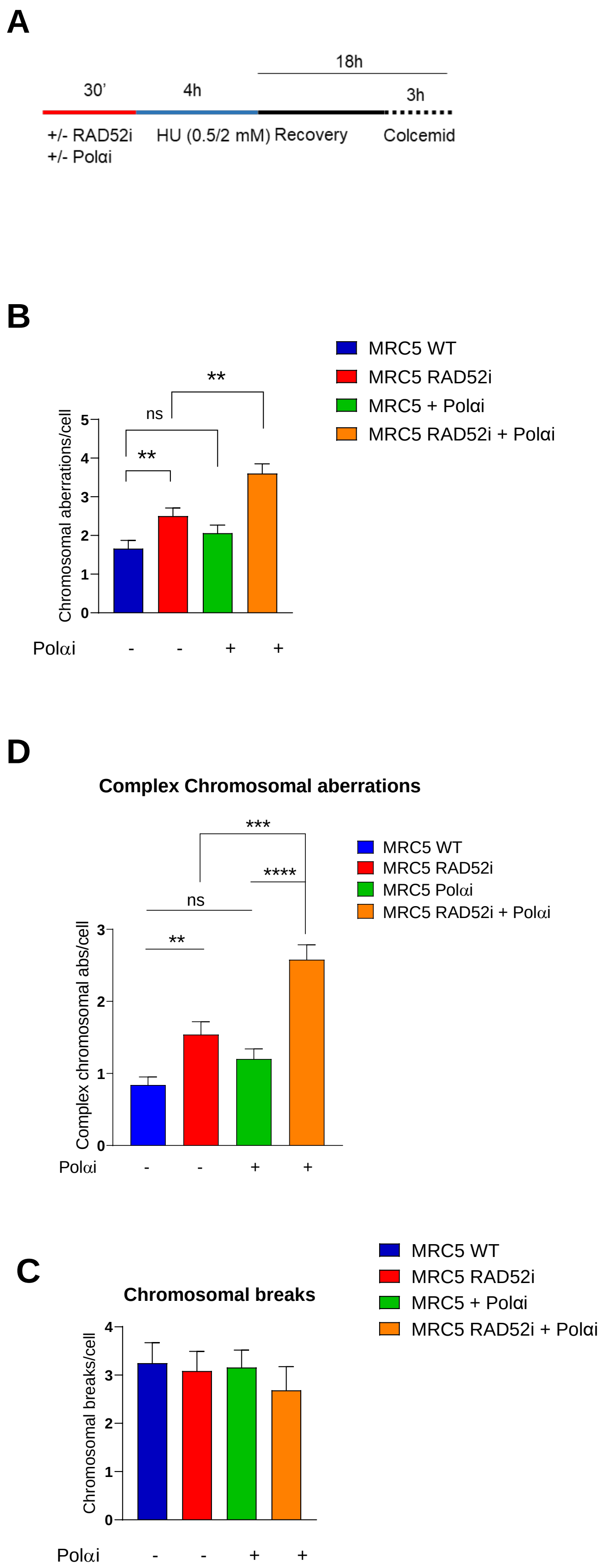


Supplementary Figure 14

### Supplementary Figure Legends

**Supplementary Figure 1. Loss of RAD52 leads to ssDNA gap detection primarily in S phase. A.** QIBC analysis of parental ssDNA exposure by immunofluorescence. Cells were treated as indicated. After the exposure to 0.5 mM hydroxyurea (HU), replication recovery was given by adding fresh medium. The ssDNA was detected in S-phase and G2-phase by IdU immunofluorescence. Cells were gated as shown in Figure 2C. **B.** Analysis of parental ssDNA in MRC5 WT and MRC5 shRAD52 cells. Graph shows the intensity of parental ssDNA (AU). Representative images are shown. **C.** Parental ssDNA exposure in U2OS treated with MRE11 inhibitor (MIRIN). Cells were treated with 0.5mM HU then released in fresh DMEM for 2-4hrs. Graph shows the mean intensity of IdU fluorescence per cells (AU). **D.** Analysis of parental ssDNA in cells after 0.5mM HU exposure. When indicated, MIRIN was added 30' before treatment. Graph shows the intensity of ssDNA per cells. In all the graphs values are from 3 different replicates and means ± SE are indicated (ns= not significant; * $P< 0.1$; ** $P<0.01$; *** $P<0.001$; **** $P<0.0001$; Kruskal-Wallis test).

**Supplementary Figure 2. RAD52 inhibition did not alter the number of S-phase cells or IdU incorporation. A.** MRC5 were subjected to EdU incorporation for 10', then were treated with HU as indicated. Graph shows the percentage of EdU-positive cells from three independent experiments. **B.** Cells were labeled with IdU and subjected to denaturing detection to assess the level of incorporation. Graph shows the mean intensity of IdU incorporation in MRC5 SV40 cells treated or not with RAD52 inhibitor and/or exposed to 2 and 0.5mM HU. **C.** Analysis of EdU-positive cells in untreated condition or after HU exposure in U2OS cells. U2OS were subjected to EdU incorporation for 10', then were treated with HU as indicated. Graph shows the percentange of EdU-positive cells from three indipendent experiments. Representative images are shown. A-C, ns= not significant, Mann-Whitney test. D, ns = not-significant, Kriuskall-Wallis test.

**Supplementary Figure 3. RAD52 inhibition did not affect fork progression.** Cells were labeled with CldU and IdU for 20' then fiber assay was performed to analyze fork progression by quantification of the IdU-labelled tracks in U2OS WT or U2OS RAD52-inhibited cells. Graph shows the IdU tract length (ns=not significant, Kruskal-Wallis test

**Supplementary Figure 4. Analysis of PrimPol-parental ssDNA association by PLA in PrimPol KO complemented cells.** Western Blot analysis of GFP-PrimPol expression. LAMIN B1 was used as a loading control. MRC5 KO PrimPol were transfected with peGFP-PrimPol WT. Twenty-four hours after transfections, cells were treated with IdU for 20 hours, released for 2 hours in fresh medium and subjected to RAD52i or MIRIN and HU (0.5 mM). Graph shows the number of PLA spots per nucleus. (ns = not significant; *P < 0.1; ****P < 0.0001; Kruskal-Wallis test).

**Supplementary Figure 5. Recruitment of Polα in S-phase is increased in RAD52 inhibited cells. A.** Analysis of Polα-parental ssDNA association by PLA. U2OS WT were treated as indicated. Graph shows the PLA spots only in EdU positive cells. **B.** Analysis of Polα-parental ssDNA in EdU negative cells. In all graphs the datapoints are from 3 independent replicates and the means ± SE are indicated (ns= not significant;***P< 0.001;****P<0.0001; Kruskal-Wallis test).

**Supplementary Figure 6. Polα recruitment in RAD52 inhibited cells does not depend on de-novo origin firing. A.** Analysis of Polα-RAD51 association by PLA in U2OS WT. Cells were treated as indicated in the scheme. PLA reaction was carried out using antibodies against IdU and Polα. The graph shows the number of PLA spot per nucleus (ns= not significant;*P < 0.1;***P< 0.001;****P<0.0001; Kruskal-Wallis test) Representative images are shown. Scale bar = 10μm.

**Supplementary Figure 7. EdU incorporation assay for the titration of Polα inhibitor** The Polαi ST1926 was tested at the concentrations reported on the graph. The cells were treated as indicated on the scheme on the left. Graph shows the mean EdU intensity. The values are presented as means ± SE (ns = not significant; *P < 0.5; **P<0.1; ***P < 0.01; ****P < 0.001; Mann–Whitney test).

**Supplementary Figure 8. RAD51 inhibition reduced the amount of parental ssDNA in RAD52 inhibited cells.** Cells were labeled with IdU for 20h, released for 2h and treated with 0.5mM HU for 4hrs. RAD51 inhibitor (BO2) was added 30' before treatment. Graph shows the amount of IdU in cells ( ns=not significant; *P<0.1; Kruskal-Wallis test).

**Supplementary Figure 9. Validation of the ΔNTD-POLA1 cell model. A.** MRC5 WT and MRC5 ΔNTD-POLA1 were transfected with siRNA directed against the 3' UTR of POLA1 and cell viability was evaluated from 24 to 72hrs post-transfection. Graph shows the percentage of cell viability at several time-points. **B.** QIBC analysis of interaction of POLA1-

RAD51 in MRC5 WT and MRC5 ΔNTD-POLA1 through PLA assay. The graph shows the number of PLA spots in RAD52 inhibited cells (n=3; ****P<0.0001; Kruskal-Wallis test).

**Supplementary Figure 10. The RAD51 T131P mutant impairs the interaction between Polα and RAD51 in the absence of RAD52.** Evaluation of RAD51-POLA1 interaction using PLA assay. Cells were transfected with RAD51 T131P and treated with 0.5mM HU for 4h. The graph shows the number of PLA spot per nucleus. (n=3; ****P < 0.0001; Kruskal-Wallis test).

**Supplementary Figure 11. Polα-primase titration experiments. A.** Polα-primase titration experiments of de novo RNA priming with products labelled with [α-32P] ATP (left panel). DNA synthesis had not observed under RNA priming buffer (middle pael) nor DNA synthesis buffer in which RNA priming is restricted (right panel). **B.** Quantification of RNA priming products in the left panel of a. RNA primer (10 nt) and reprimed RNA primer (20nt) are formed in the condition. **C.** Model explaining Polα-primase RNA primer extension assay. 5' end Cy3-dye labeled rA15 RNA primer is extended by Polα-primase DNA polymerase activity in DNA synthesis buffer and the presence of dATP. **D.** Polα-primase titration experiments on 5'-Cy3-labeled poly-rA15 primer extension. Size markers are synthesized Cy3-labelled poly(dT) ssDNA.

**Supplementary Figure 12. Analysis of RAD51-stimulated Polα priming using long circular ssDNA φX174 Virion DNA as a template.** Polα/Primase no longer generates 70 nt RNA product in the presence of RAD51 but produces extra-long primer. The 5'-Cy3-labeled poly-rA15 primer was extended by DNA synthesis activity of Polα/Primase in the presence of dATP and on RAD51 **(A-C)** or RPA **(D-F)** titration. Size markers are synthesized Cy3-labelled poly(dT) ssDNA.

**Supplementary Figure 13. Polα-primase titration experiments with mutant RAD51 proteins. A.** RAD51-T131P and RAD51-F86E titration on Polα-primase de novo RNA synthesis on dT70 mer templates, with products labelled with [α-32P]ATP. **B.** Quantification of RNA products in the experiment directly above. 10 and 20 nt RNA priming product (grey), template size RNA product (green) and long RNA product that is stacked in the well (red). **C.** RPA was incubated with poly(dT)70 (500 nM, molecules), and the RPA–DNA complexes were analyzed by EMSA. The mobility of the ssDNA–RPA complex was shifted in several steps. Poly(dT)70 was saturated at approximately 3 molecules of RPA per DNA molecule. These results indicate that three RPA molecules bound to 70 nt ssDNA and are consistent with the previously reported binding site size of RPA (20-30 nt per RPA). **D.** RAD51 binds poly(dT)70. 500 nM of each DNA substrate were incubated with indicated amount of RAD51. The bound and free DNA molecules were resolved by EMSA and quantified (DNA density counts of lane 1 in each panel are defined as 100%).

**Supplementary Figure 14. Combined treatment with RAD52i and POLA1i, increases total chromosomal damage, fusions and exchanges.** Analysis of chromosomal aberrations in Polα inhibited cells. **A. Schematic representation of treatments. B.** Analysis of total chromosomal aberrations in MRC5SV40 cells treated as in the experimental scheme. The graphs show the means of total chromosomal aberrations per cell and the means of chromosomal breaks (**C**) and complex chromosomal aberrations per cell (**D**). In all the graphs the results are from three independent experiments and are presented as means ± SE (ns = not significant; *P < 0.1; **P<0.1; ***P < 0.001; ****P < 0.0001; ANOVA test).
